# Phylogenomics reveals the relationships of butterflies and moths (Lepidoptera): providing the potential landscape using universal single copy orthologues

**DOI:** 10.1101/2022.10.14.512238

**Authors:** Qi Chen, Min Deng, Wei Wang, Xing Wang, Liu-Sheng Chen, Guo-Hua Huang

## Abstract

**Background:** A robust and stable phylogenetic framework is a fundamental goal of evolutionary biology. As the third largest insect order following by Diptera and Coleoptera in the world, lepidoptera (butterflies and moths) play a central role in almost every terrestrial ecosystem as the indicators of environmental change and serve as important models for biologists exploring questions related to ecology and evolutionary biology. However, for such charismatic insect group, the higher-level phylogenetic relationships among its superfamilies are still poorly unresolved.

**Results:** we increased taxon sampling among Lepidoptera (40 superfamilies and 76 families contained 286 taxa) and filtered the unqualified samples, then acquired a series of large amino-acid datasets from 69,680 to 400,330 for phylogenomic reconstructions. Using these datasets, we explored the effect of different taxon sampling on tree topology by considering a series of systematic errors using ML and BI methods. Moreover, we also tested the effectiveness in topology robustness among the three ML-based models. The results showed that taxon sampling is an important determinant in tree robustness of accurate lepidopteran phylogenetic estimation. Long-branch attraction (LBA) caused by site-wise heterogeneity is a significant source of bias given rise to topologies divergence of ditrysia in phylogenomic reconstruction. Phylogenetic inference showed a most comprehensive framework by far to reveal the relationships among lepidopteran superfamilies, but limited by taxon sampling, it could only represent the current understanding of the lepidopteran tree of life. The relationships within the species-rich and relatively rapid radiation Ditrysia and especially Apoditrysia remain poorly unresolved, which need to increase taxon sampling and adopt lineage-specific genes for further phylogenomic reconstruction.

**Conclusions:** The present study further expands the taxon sampling of lepidopteran phylogeny and provides a potential phylogenomic foundation for further understanding its current higher-level relationships.

## Background

A robust and stable phylogenetic framework is a fundamental goal of evolutionary biology [1–3], which could effectively help evolutionary biologists for inferring the origin and evolution of species, detecting molecular adaptation, understanding morphological character evolution and reconstructing demographic changes in certain taxa [4, 5]. Traditional molecular systematics is dominated by Sanger sequencing, phylogenetic reconstruction depends on data from one or a few genes generated typically using PCR amplification [6, 7]. However, the phylogenetic relationships for some lineages are controversial by using this method due to the limitation of insufficient phylogenetic signal with an inaccurate tree [8–11]. With the emergence of new sequencing technologies, next-generation sequencing (NGS) with genome-scale data provides an opportunity to address such long-standing controversies across all lineages of the tree of life [12–14].

Many strategies have been developed and driven by this trend to the forefront in practice. Representative technologies employed in deep phylogenetics include phylotranscriptomics [15], anchored hybrid enrichment (AHE) [16], ultraconserved element enrichment (UCE) [17], and automated target restricted assembly method (aTRAM) [18], which have been proved be useful for constructing robust phylogenies and successfully employed to address a wide variety of questions for some lineages [18–20]. However, the limitations of these technologies with specific materials and time-consuming data calculation are still obvious undeniably on a trial, making it more boundedness in a deep phylogeny in many cases [21, 22]. The continuing decrease of sequencing costs in recent years have made low-coverage whole-genome sequencing (LCWGS) more economically feasible, and WGS data are becoming widely applied in ecological and evolutionary studies [23, 24]. Benchmarking universal single-copy orthologs (BUSCOs) are a set of conservative genes of universal single-copy orthologs identified using OrthoDB [25], which have been commonly used for quantitative assessment of genome assembly [26, 27]. A series of recent studies based on LCWGS data turned out these genes are reliable and reproducible markers in higher phylogeny of Collembola [28], green lacewings [29], and elongate-bodied springtails [30]. Nevertheless, for most conspicuous and phylogenetically controversial lineages, such as lepidoptera, there is an insufficient practice to deliver the higher-level phylogenomics needs via using these markers on the basic of relatively abundant available genomic resources, in that way, how effectiveness of that for complicated lineages is unknown.

Lepidoptera (butterflies and moths) is the third largest order following by Diptera and Coleoptera in the world, which consists with more than 157,000 described species belonging to 137 families within 43 superfamilies [31]. Within the order, there are many damaging pests (e.g. *Bombyx*, *Cydia*, *Helicoverpa*, *Manduca*, *Spodoptera*) in agriculture and most important model organisms for exploring questions related to ecology and evolutionary biology [32]. As herbivores, pollinators and prey, lepidopteran insects widely exist in nearly all continents and a variety of habitats [33], and play a central role in the study of speciation, community ecology, biogeography, and climate change [34, 35]. And the intricate relationships with their host plants have long provided an ideal model for evolutionary biologists to investigating coevolutionary dynamics of plant-insect [36, 37]. However, for such charismatic insect groups, the phylogenetics of some major lepidopteran groups have not robustly resolved in higher-level relationships [35, 38], especially for that of the largest radiation and species-rich lineages Ditrysia, which potentially constrains our understanding of the reason of species diversification and morphological evolution of key clades of lepidoptera.

Previous deep-level molecular systematics for lepidoptera mainly focus on a small number of genes obtained via traditional Sanger sequencing [38–42] or complete mitochondrial genomes generated by using next generation-sequencing methods [43, 44]. Nonetheless, unambiguous position among several superfamilies in lepidoptera, especially within ditrysia, remain uncertain due to the poor phylogenetic signal of loci and/or limited taxon sampling in these researches (Fig. 1A-1B). Recent several phylogenomic studies based on transcriptomes and target enrichment approaches have greatly expanded the numbers of loci (from 741 to 2,948 genes) or taxa used to address deep phylogenetic relationships within lepidoptera [35, 45–48], which provide many of the surprising results that obtained in previous Sanger sequencing based studies. However, for the key clades (e.g. Apoditrysia or Eulepidoptera), the phylogenetic backbone among their superfamilies have been still unsettled due to the conflicting evidence of crucial nodes each other and inconsistent taxon sampling among these studies, which hinder exceedingly our ability to understand the early lepidopteran diversification (Fig. 1C-1E). A recent study based on 34 superfamilies and 72 families of lepidoptera using 331 protein-coding genes assesses the potential causes and consequences of the conflicting phylogenetic hypotheses and presents a comprehensive phylogenetic framework of butterflies and moths [49]. However, in common with other studies, limited by taxon sampling, the positions among superfamilies within Apoditrysia are still uncertainty and need further extra attention (Fig. 1F). Therefore, increasing taxon sampling continuously to reconstruct a stable and well-supported phylogenomic tree among key clades might be crucial at present for discussion on obscure evolutionary relationships of Lepidoptera.

**Figure 1.**
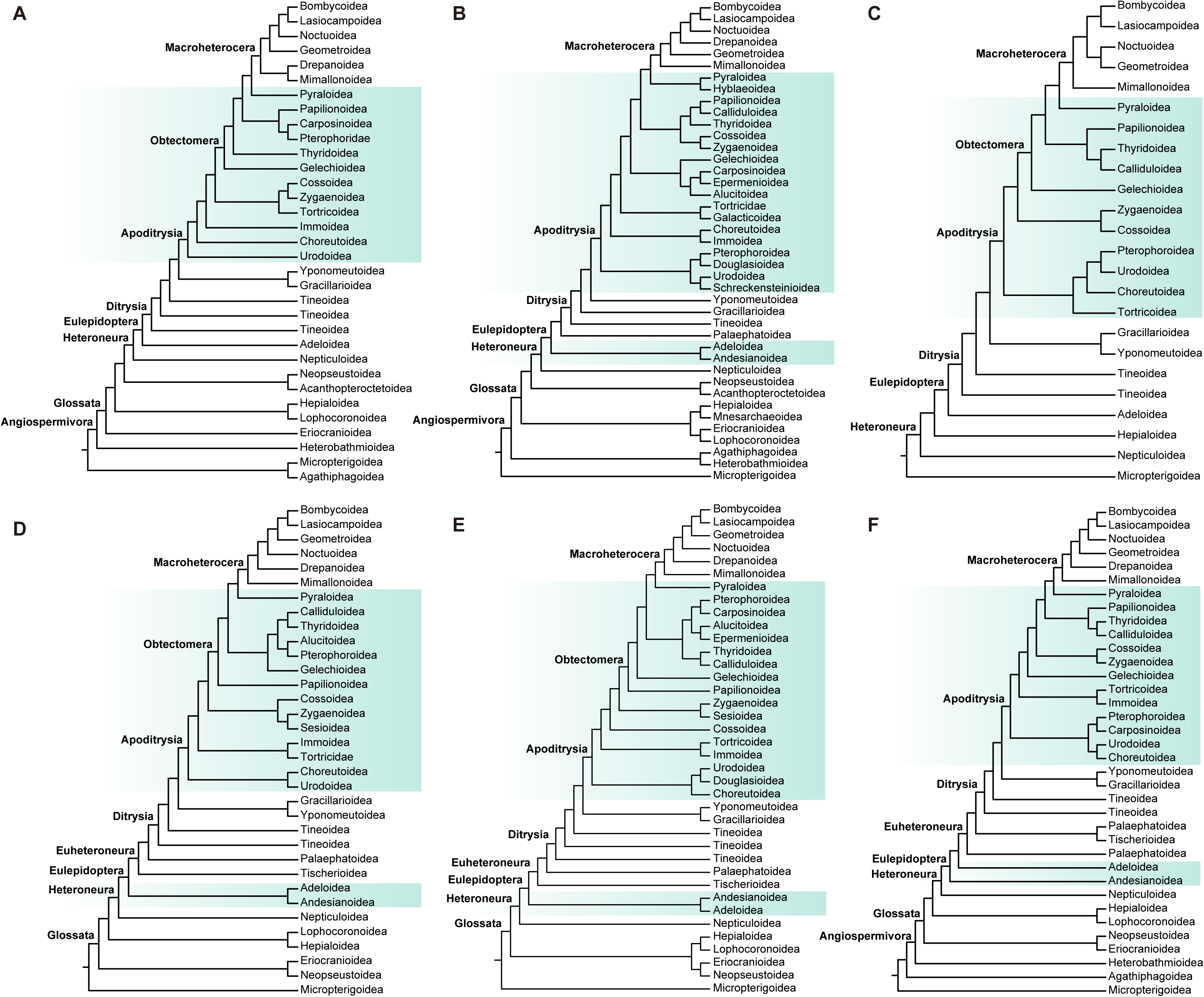
Previous hypotheses of deep-level relationships in Lepidoptera. **(A)** Maximum likelihood tree inferred from 19 protein-coding nuclear genes and 483 taxa by Regier et al. (2013) [39]. **(B)** Maximum likelihood tree based on morphological characters combined with one mitochondrial and seven nuclear protein-coding genes, as well as 473 exemplar species by Heikkilä et al. (2015) [40]. **(C)** Maximum likelihood tree estimated from AHE and transcriptomic data of 76 exemplar species in 42 families by Breinholt et al. (2018) [47]; **(D)** Evolutionary tree from a maximum-likelihood analysis of 749,791 amino acid sites from transcriptomes of 186 species by Kawahara et al. (2019) [35]. **(E)** Phylogenetic tree inferred using 1,835 CDS nucleotides of 172 taxa by Mayer et al. (2021) [48]. **(F)** Maximum likelihood tree estimated using 331 protein-coding genes of 200 taxa representing 83% of lepidopteran superfamilies by Rota et al. (2022) [49]. Gradient colors indicate the complicated phylogenetic relationships among lepidopteran superfamilies across the different studies.

To infer the phylogenetic relationships within Lepidoptera, we increased taxon sampling among Lepidoptera (40 superfamilies and 76 families contained 286 taxa) and filtered the unqualified samples, then acquired a series of large phylogenomic amino-acid datasets derived from genomes and transcriptomes against the insecta (n=1,367) and lepidoptera (n=5,286) reference gene sets. Using these datasets, we explored the effect of different taxon sampling for tree topology robustness over considering systematic errors. Simultaneously, we also compared the differences in phylogenetics tree reconstructed among the three models under the maximum likelihood method. The present study would provide a potential phylogenomic foundation for evaluating hypotheses on higher-level relationships within lepidoptera.

## Results

### Genome and transcriptome assembly and BUSCO assessment

Illumina sequencing yielded approximately 280G raw data, which coverage ranged from 13.16 × to 82.80 × for the 53 newly sequenced lepidopteran genomes (Table 1). *de novo* genome assembly was performed using Spades, assembly statistics were summarized in Table 1, ranging from: genome size of 244.49-608.19 Mb, 32,071-137,263 scaffolds, N50 length of 4.642-29.888 kb, longest scaffold of 46.388-1060 kb and 32.31%-40.15% GC content. BUSCO completeness against the Insecta (n=1,367) and lepidoptera (n=5,286) reference set were 52.7% (720)-94.1% (1286) and 42.0% (2,220)-93.5% (4,947) complete BUSCOs, respectively. Furthermore, a series of lepidopteran genomes or transcriptomes (233 taxa) were downloaded from NCBI to expand our taxon sampling. For genomes, the vast majority of them had been assembled at the chromosome level. We collected the assembled information of them again and summarized in Table S2 (Additional file 1: Table S2), ranging from: genome size of 229.694-1305.982 Mb, 19-293,367scaffolds, N50 length of 0.028-95.393 Mb, longest scaffold of 0.02-387.839 Mb and 31.6%-39.76% GC content. BUSCO completeness against the Insecta (n=1,367) and lepidoptera (n=5,286) reference set were 45.3% (620)-99% (1,353) and 46.2% (2444)-99.3% (5,250) complete BUSCOs, respectively. For transcriptomes, Trinity was employed for *de novo* assembly. Assembled statistics were assessed and shown in Table S3 (Additional file 1: Table S3), ranging from: transcriptome size of 13.464-225.937 Mb, 14,895-458,022 scaffolds, N50 length of 0.257-2.714 kb, longest scaffold of 5.986-54.961 kb and 33.49%-44.06% GC content. BUSCO completeness against the Insecta (n=1367) and lepidoptera (n=5286) reference set were 30.6% (418)-95.1% (1,301) and 19.2% (1,017)-89.4% (4,727) complete BUSCOs, respectively. However, the completeness of some transcriptomes was below 30% (completeness from 2.4% to 29.5%) against the Insecta (n=1,367) gene set (Additional file 1: Table S4), we had to exclude those taxa with lower completeness to ensure adequate phylogenetic signals. Finally, 263 taxa representing 38 of 41 currently recognized lepidopteran superfamilies were adopted for following phylogenetic inference.

**Table 1.** The statistics for genome assembly of specimens used for phylogenomic analysis in the present study.

| Superfamily | Family | Species | Average<br>depth<br>(×) | Total<br>length<br>(Mb) | Scaffold<br>number | N50<br>length<br>(kb) | Longest<br>scaffold<br>(kb) | GC% | Complete BUSCOs (Insecta: n=1367) |  |  |  |  | Complete BUSCOs (lepidoptera: n=5286) |  |  |  |  | SRA number |
| --- | --- | --- | --- | --- | --- | --- | --- | --- | --- | --- | --- | --- | --- | --- | --- | --- | --- | --- | --- |
| Adeloidea | Adelidae | *Nemophora* sp1. | 19.12× | 387.96 | 84272 | 16.493 | 70.72 | 37.20 | C:898 | S:895 | D:3 | F:321 | M:148 | C:3419 | S:3377 | D:42 | F:636 | M:1231 | XXX |
| Adeloidea | Adelidae | *Nemaphora* sp2. | 28.39× | 400.16 | 124504 | 26.582 | 55.907 | 38.18 | C:876 | S:862 | D:14 | F:339 | M:152 | C:3359 | S:3266 | D:93 | F:668 | M:1259 | XXX |
| Adeloidea | Adelidae | *Nemaphora* sp3. | 29.02× | 562.92 | 131601 | 25.283 | 59.822 | 36.76 | C:843 | S:836 | D:7 | F:345 | M:179 | C:3172 | S:3096 | D:76 | F:682 | M:1432 | XXX |
| Adeloidea | Heliozelidae | *Antispila* sp. | 30.32× | 244.49 | 44303 | 7.347 | 95.705 | 40.15 | C:979 | S:974 | D:5 | F:259 | M:129 | C:3479 | S:3423 | D:56 | F:511 | M:1296 | XXX |
| Adeloidea | Heliozelidae | *Heliozela* sp. | 34.84× | 377.58 | 68332 | 12.49 | 86.205 | 39.69 | C:916 | S:909 | D:7 | F:288 | M:163 | C:3312 | S:3268 | D:44 | F:500 | M:1474 | XXX |
| Bombycoidea | Bombycidae | *Gunda sesostris* | 14.73× | 385.93 | 95788 | 16.455 | 185.474 | 38.56 | C:1051 | S:1046 | D:5 | F:201 | M:115 | C:4229 | S:4194 | D:35 | F:551 | M:506 | XXX |
| Bombycoidea | Bombycidae | *Penicillifera tamsi* | 24.38× | 307.88 | 38473 | 6.643 | 228.977 | 34.96 | C:1188 | S:1182 | D:6 | F:118 | M:61 | C:4736 | S:4684 | D:52 | F:342 | M:208 | XXX |
| Bombycoidea | Bombycidae | *Trilocha varians* | 18.59× | 265.66 | 97139 | 22.301 | 423.579 | 35.77 | C:1022 | S:1003 | D:19 | F:191 | M:154 | C:3716 | S:3631 | D:85 | F:714 | M:856 | XXX |
| Bombycoidea | Bombycidae | *Penicillifera lactea* | 18.57× | 358.10 | 81485 | 13.976 | 90.817 | 35.33 | C:1108 | S:1100 | D:8 | F:176 | M:83 | C:4477 | S:4399 | D:78 | F:457 | M:352 | XXX |
| Bombycoidea | Endromidae | *Andraca olivacea* | 19.75× | 349.72 | 74935 | 12.532 | 391.074 | 37.61 | C:1186 | S:1174 | D:12 | F:121 | M:60 | C:4702 | S:4550 | D:152 | F:364 | M:220 | XXX |
| Bombycoidea | Endromidae | *Andraca trilochoides* | 37.43× | 385.43 | 93654 | 14.544 | 498.114 | 40.11 | C:1161 | S:1138 | D:23 | F:126 | M:80 | C:4572 | S:4442 | D:130 | F:465 | M:249 | XXX |
| Bombycoidea | Endromidae | *Comparmustilia semiravida* | 22.70× | 358.25 | 50078 | 6.141 | 406.029 | 37.07 | C:1230 | S:1216 | D:14 | F:64 | M:73 | C:4827 | S:4730 | D:97 | F:286 | M:173 | XXX |
| Bombycoidea | Endromidae | *Comparmustilia semiravida* | 38.91× | 350.30 | 51680 | 6.714 | 296.862 | 36.26 | C:1208 | S:1206 | D:2 | F:103 | M:56 | C:4817 | S:4758 | D:59 | F:286 | M:183 | XXX |
| Bombycoidea | Endromidae | *Dalailama vadim* | 18.32× | 367.07 | 48384 | 7.038 | 1026 | 39.62 | C:1231 | S:1127 | D:104 | F:62 | M:74 | C:4722 | S:4621 | D:101 | F:344 | M:220 | XXX |
| Bombycoidea | Endromidae | *Endromis*<br>*versicolora* | 19.12× | 364.63 | 43644 | 7.879 | 103.702 | 38.12 | C:1130 | S:1128 | D:2 | F:145 | M:92 | C:4574 | S:4524 | D:50 | F:435 | M:277 | XXX |
| Bombycoidea | Endromidae | *Mustilia*<br>*falcipennis* | 18.22× | 391.13 | 78002 | 13.191 | 185.636 | 36.02 | C:1026 | S:1022 | D:4 | F:219 | M:122 | C:4184 | S:4114 | D:70 | F:609 | M:493 | XXX |
| Bombycoidea | Endromidae | *Mustilizans*<br>*capella* | 21.31× | 332.59 | 39586 | 4.877 | 1060 | 38.06 | C:1286 | S:1267 | D:19 | F:39 | M:42 | C:4938 | S:4855 | D:83 | F:237 | M:111 | XXX |
| Bombycoidea | Endromidae | *Mustilizans*<br>*shennongi* | 25.72× | 287.32 | 37950 | 5.85 | 201.839 | 35.31 | C:1257 | S:1252 | D:5 | F:69 | M:41 | C:4947 | S:4871 | D:76 | F:226 | M:113 | XXX |
| Bombycoidea | Endromidae | *Oberthueria*<br>*formosibia* | 17.11× | 408.42 | 81149 | 13.277 | 202.905 | 36.54 | C:1169 | S:1152 | D:17 | F:138 | M:60 | C:4697 | S:4545 | D:152 | F:386 | M:203 | XXX |
| Bombycoidea | Endromidae | *Oberthueria*<br>*falcigera* | 17.82× | 327.57 | 39448 | 5.98 | 140.413 | 36.56 | C:1237 | S:1233 | D:4 | F:72 | M:58 | C:4876 | S:4816 | D:60 | F:271 | M:139 | XXX |
| Bombycoidea | Endromidae | *Oberthueria* sp. | 25.13× | 480.30 | 137263 | 27.895 | 204.677 | 36.61 | C:1047 | S:1002 | D:45 | F:184 | M:136 | C:4193 | S:3913 | D:280 | F:671 | M:422 | XXX |
| Bombycoidea | Endromidae | *Oberthueria*<br>*caeca* | 34.15× | 370.51 | 59621 | 6.387 | 171.903 | 36.49 | C:1191 | S:1171 | D:20 | F:114 | M:62 | C:4793 | S:4676 | D:117 | F:307 | M:186 | XXX |
| Bombycoidea | Endromidae | *Prismosticta*<br>*microprisma* | 27.60× | 451.28 | 86632 | 14.012 | 248.568 | 35.76 | C:1134 | S:1120 | D:14 | F:158 | M:75 | C:4581 | S:4419 | D:162 | F:460 | M:245 | XXX |
| Bombycoidea | Endromidae | *Prismosticta*<br>*regalis* | 31.47× | 414.60 | 94059 | 16.921 | 106.428 | 35.95 | C:1059 | S:1053 | D:6 | F:212 | M:96 | C:4349 | S:4226 | D:123 | F:569 | M:368 | XXX |
| Bombycoidea | Endromidae | *Promustilia*<br>*yajiangensis* | 18.74× | 295.99 | 51193 | 8.962 | 614.309 | 36.48 | C:1199 | S:1181 | D:18 | F:84 | M:84 | C:4607 | S:4547 | D:60 | F:400 | M:279 | XXX |
| Bombycoidea | Endromidae | *Pseudandraca*<br>*flavamaculata* | 21.49× | 400.00 | 101503 | 19.098 | 121.522 | 36.44 | C:1124 | S:1102 | D:22 | F:159 | M:84 | C:4527 | S:4354 | D:173 | F:486 | M:273 | XXX |
| Bombycoidea | Endromidae | *Smerkata fusca* | 18.54× | 338.13 | 57226 | 8.91 | 102.884 | 36.04 | C:1111 | S:1108 | D:3 | F:172 | M:84 | C:4443 | S:4385 | D:58 | F:478 | M:365 | XXX |
| Eriocranioidea | Eriocraniidae | *Eriocrania*<br>*komaii* | 25.63× | 490.50 | 82076 | 14.15 | 98.328 | 34.21 | C:1091 | S:1082 | D:9 | F:177 | M:99 | C:3519 | S:3466 | D:53 | F:351 | M:1416 | XXX |
| Gelechioidea | Gelechiidae | *Monochroa* sp. | 18.10× | 425.10 | 56103 | 9.082 | 185.076 | 37.30 | C:1097 | S:1092 | D:5 | F:171 | M:99 | C:4403 | S:4351 | D:52 | F:448 | M:435 | XXX |
| Geometroidea | Geometridae | *Hydrelia*<br>*parvulata* | 16.15× | 362.33 | 80127 | 14.997 | 104.788 | 37.14 | C:1062 | S:1053 | D:9 | F:197 | M:108 | C:4102 | S:4044 | D:58 | F:576 | M:608 | XXX |
| Micropterigoidea | Micropterigidae | *Vietomartyria*<br>*maoershana* | 19.01× | 356.06 | 62508 | 9.477 | 112.568 | 33.00 | C:1023 | S:1016 | D:7 | F:230 | M:114 | C:3183 | S:3134 | D:49 | F:363 | M:1740 | XXX |
| Micropterigoidea | Micropterigidae | *Vietomartyria*<br>*wuyunjiena* | 21.50× | 409.17 | 56712 | 8.681 | 134.675 | 32.37 | C:1062 | S:1053 | D:9 | F:181 | M:124 | C:3147 | S:3090 | D:57 | F:364 | M:1775 | XXX |
| Micropterigoidea | Micropterigidae | *Vietomartyria*<br>*wuyunjiena* | 17.35× | 398.51 | 58702 | 9.229 | 128.505 | 32.34 | C:1039 | S:1029 | D:10 | F:206 | M:122 | C:3056 | S:3009 | D:47 | F:360 | M:1870 | XXX |
| Micropterigoidea | Micropterigidae | *Vietomartyria* sp. | 18.95× | 388.77 | 32071 | 4.642 | 274.185 | 32.31 | C:1165 | S:1153 | D:12 | F:110 | M:92 | C:3488 | S:3432 | D:56 | F:298 | M:1500 | XXX |
| Micropterigoidea | Micropterigidae | *Neomicropteryx*<br>*bifurca* | 26.14× | 455.29 | 47882 | 7.451 | 208.592 | 33.04 | C:1135 | S:1122 | D:13 | F:137 | M:95 | C:3447 | S:3400 | D:47 | F:276 | M:1563 | XXX |
| Neopseustoidea | Neopseustidae | *Neopseustis*<br>*fanjingshana* | 23.94× | 395.04 | 73568 | 13.779 | 95.94 | 34.45 | C:952 | S:946 | D:6 | F:259 | M:156 | C:3126 | S:3058 | D:68 | F:480 | M:1680 | XXX |
| Neopseustoidea | Neopseustidae | *Neopseustis*<br>*rectagnathos* | 20.11× | 410.55 | 127082 | 27.763 | 109.857 | 34.29 | C:765 | S:757 | D:8 | F:363 | M:239 | C:2552 | S:2484 | D:68 | F:571 | M:2163 | XXX |
| Nepticuloidea | Nepticulidae | Nepticulidae sp. | 21.87× | 309.32 | 118589 | 29.888 | 55.553 | 32.35 | C:742 | S:737 | D:5 | F:329 | M:296 | C:2220 | S:2174 | D:46 | F:489 | M:2577 | XXX |
| Nepticuloidea | Nepticulidae | *Ectoedemia* sp. | 19.99× | 443.18 | 88042 | 15.312 | 908.378 | 36.49 | C:844 | S:821 | D:23 | F:261 | M:262 | C:2777 | S:2692 | D:85 | F:489 | M:2020 | XXX |
| Papilionoidea | Lycaenidae | *Cyaniris*<br>*semiargus* | 28.44× | 378.05 | 104186 | 23.579 | 46.388 | 36.42 | C:987 | S:982 | D:5 | F:250 | M:130 | C:3876 | S:3840 | D:36 | F:695 | M:715 | XXX |
| Papilionoidea | Lycaenidae | *Cyaniris*<br>*semiargus* | 82.80× | 412.88 | 96900 | 18.913 | 229.547 | 37.69 | C:1048 | S:1034 | D:14 | F:202 | M:117 | C:4058 | S:3978 | D:80 | F:540 | M:688 | XXX |
| Papilionoidea | Lycaenidae | *Glaucopsyche*<br>*alexis* | 32.94× | 429.82 | 110463 | 23.868 | 54.331 | 35.90 | C:942 | S:940 | D:2 | F:282 | M:143 | C:3835 | S:3786 | D:49 | F:718 | M:733 | XXX |
| Papilionoidea | Lycaenidae | *Maculinea* sp. | 36.60× | 451.10 | 60646 | 10.652 | 115.101 | 36.12 | C:1150 | S:1146 | D:4 | F:130 | M:87 | C:4480 | S:4438 | D:42 | F:369 | M:437 | XXX |
| Papilionoidea | Lycaenidae | *Phengaris teleius* | 38.76× | 450.38 | 52520 | 8.807 | 132.853 | 35.78 | C:1179 | S:1176 | D:3 | F:115 | M:73 | C:4601 | S:4547 | D:54 | F:359 | M:326 | XXX |
| Papilionoidea | Lycaenidae | *Phengaris teleius* | 37.79× | 452.89 | 54658 | 8.991 | 132.288 | 35.77 | C:1169 | S:1163 | D:6 | F:116 | M:82 | C:4597 | S:4545 | D:52 | F:352 | M:337 | XXX |
| Papilionoidea | Lycaenidae | *Phengaris alcon* | 27.12× | 436.65 | 58669 | 10.556 | 162.504 | 35.72 | C:1126 | S:1122 | D:4 | F:156 | M:85 | C:4413 | S:4376 | D:37 | F:414 | M:459 | XXX |
| Papilionoidea | Nymphalidae | *Lethe nigrifascia* | 19.52× | 409.31 | 63488 | 10.998 | 150.778 | 36.13 | C:1110 | S:1107 | D:3 | F:160 | M:97 | C:4472 | S:4404 | D:68 | F:452 | M:362 | XXX |
| Papilionoidea | Papilionidae | *Luehdorfia chinensis* | 17.25× | 608.19 | 106600 | 20.067 | 89.676 | 37.34 | C:974 | S:967 | D:7 | F:250 | M:143 | C:4059 | S:3998 | D:61 | F:709 | M:518 | XXX |
| Pyraloidea | Pyralidae | *Pyralidae* sp. | 16.07× | 337.01 | 46613 | 8.195 | 117.666 | 37.69 | C:1177 | S:1174 | D:3 | F:116 | M:74 | C:4618 | S:4574 | D:44 | F:369 | M:299 | XXX |
| Gracillarioidea | Gracillariidae | *Caloptilia aceris* | 21.57× | 386.92 | 116875 | 25.398 | 68.664 | 38.24 | C:1021 | S:1000 | D:21 | F:236 | M:110 | C:4031 | S:3823 | D:208 | F:612 | M:643 | XXX |
| Tischerioidea | Tischeriidae | *Tischeria decidua* | 24.83× | 498.54 | 91124 | 16.588 | 110.527 | 36.94 | C:937 | S:929 | D:8 | F:262 | M:168 | C:3411 | S:3366 | D:45 | F:489 | M:1386 | XXX |
| Tischerioidea | Tischeriidae | *Coptotriche* sp. | 13.16× | 469.96 | 128984 | 28.122 | 126.08 | 33.89 | C:720 | S:713 | D:7 | F:357 | M:290 | C:2546 | S:2501 | D:45 | F:703 | M:2037 | XXX |

### Data and ML model effectiveness

To explore the phylogenetic relationships among lepidopteran superfamilies and assess the effectiveness of various taxon sampling for tree topology robustness, we conducted phylogenetic reconstructions using a series of datasets with supplementing diverse analytical methods to reduce the possible systematic errors. For insecta datasets, number of amino acid sites were ranged from 69,680 to 337,588, contained loci from 248 to 1,004 with average locus length of 280.97-336.24, respectively (Table 2). Furthermore, relatively sufficient parsimony informative (65.568-67.796) and few missing (4.867-5.787) sites indicated that these datasets had some specific phylogenetic signal for tree topology reconstructions to a certain extent. However, there was an obvious insufficiency with limited loci and shorter sites in length as well as the distant lineages among universal single-copy orthologous for them due to the objective limitation of reference Insecta BUSCO gene sets (n=1,367). We could not be sure that these datasets were suitable for revealing phylogenetic relationships of lepidopteran superfamilies. We supplemented with three lepidoptera datasets to verify the above tree topology robustness. For these three datasets, number of amino acid sites were ranged from 234,774 to 400,330, included loci from 997 to 1,266 with mean locus length of 235.48-316.22, respectively (Table 2).

**Table 2.** An overview of BUSCO amino acid matrices used for phylogenetic inference.

| **Matrix** | **Number**<br>**of taxa** | **Lineages dataset** | **BUSCO completeness of taxa** | **Number**<br>**of loci** | **Number**<br>**of sites** | **Missing**<br>**sites (%)** | **Parsimony**<br>**informative sites (%)** | **Average**<br>**locus length** |
| --- | --- | --- | --- | --- | --- | --- | --- | --- |
| BUSCO30 | 268 | Insecta (n=1367) | 30.6% (418) ~ 99% (1353) | 248 | 69,680 | 5.787 | 67.349 | 280.97 |
| BUSCO60 | 241 | Insecta (n=1367) | 60.1% (822) ~ 99% (1353) | 640 | 186,639 | 5.605 | 67.717 | 291.62 |
| BUSCO70 | 224 | Insecta (n=1367) | 71% (971) ~ 99% (1353) | 784 | 239,127 | 5.532 | 67.796 | 305.01 |
| BUSCO80 | 187 | Insecta (n=1367) | 80.3% (1097) ~ 99% (1353) | 1004 | 337,588 | 4.867 | 65.568 | 336.24 |
| BUSCO30_test | 250 | Insecta (n=1367) | 48.8% (666) ~ 99% (1353) | 248 | 69,680 | 5.787 | 67.349 | 280.97 |
| lep_busco | 250 | lepidoptera (n=5286) | 33.3% (1761) ~ 99.3% (5250) | 1017 | 289,668 | 6.883 | 71.358 | 284.83 |
| noditrysia_busco | 41 | lepidoptera (n=5286) | 33.3% (1761) ~ 88.8% (4697) | 997 | 234,774 | 16.692 | 51.637 | 235.48 |
| ditrysia_busco | 208 | lepidoptera (n=5286) | 43.2% (2286) ~ 99.3% (5250) | 1266 | 400,330 | 4.958 | 68.172 | 316.22 |

All ML analyses using these datasets yielded 22 phylogenetic trees with highly ultrafast bootstrap supports for most branches (Additional file 2: Figure S1-S22). Average ultrafast bootstraps (aUBF) were ranged from partition scheme of 97.86-100, GHOST model of 99.28-100, and PMSF model of 98.35-100, respectively (Additional file 1: Table S5). Moreover, all phylogenetic reconstructions were clearly shown that the backbone of nonditrysian superfamilies were almost consistent compared to the tree topologies each other (Fig. 2, Additional file 1: Table S6, Additional file 2: Figure S1-S30), and the present result is also observed in previous reports (Fig. 1) [30, 35, 39, 40, 46, 47, 49].

**Figure 2.**
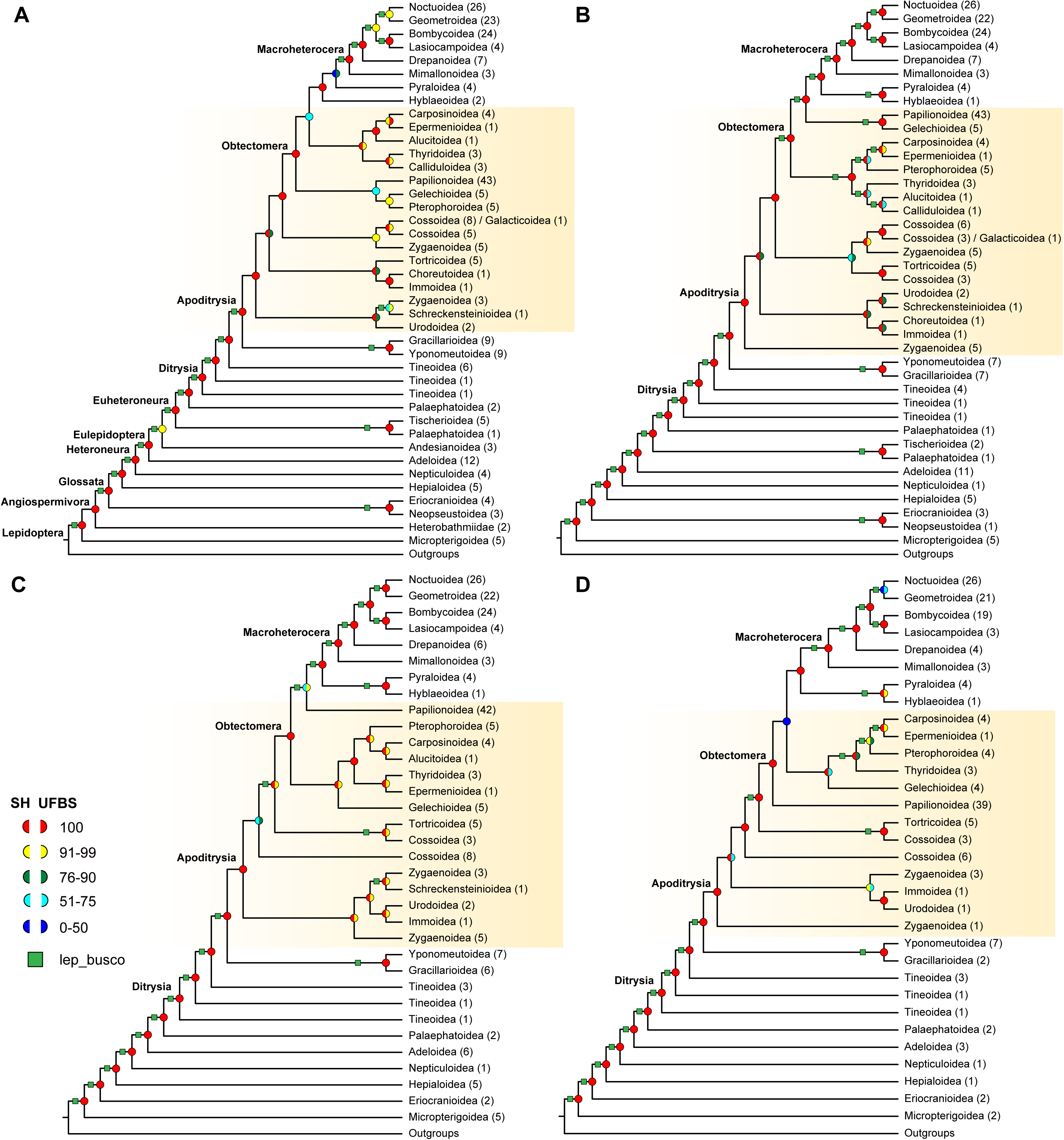
An overview of deep-level relationships among Lepidoptera created from PMSF model based on maximum likelihood analyses of different insecta datasets. (A) BUSCO30; (B) BUSCO60; (C) BUSCO70; (D) BUSCO80. Full dataset descriptions are available in Table 2. The topology is similar to that of lep_busco is emphasized on the branch by using green square. Numbers after the labels of superfamily show the total number of taxa for this superfamily in corresponding dataset. Gradient colors indicate the complicated phylogenetic relationships among lepidopteran superfamilies across the different datasets and colored labels at the nodes represent the summarized status of bootstrap (SH-aLRT and UFBS) values.

However, for the species-rich lepidopteran clade Apoditrysia, phylogenetic inferences showed a variety of inconsistent tree topologies among its superfamilies (Fig. 2, Additional file 1: Table S6, Additional file 2: Figure S1-S30), it was hard to determine which topology in a certain model was most robust with a good performance in ML analyses. We carried out topologies test for the resultant three alternative topologies with the corresponding dataset in IQtree. Not surprised, the computing resource-consuming model PMSF presented almost maximum likelihood values with relatively high *p*-values in hypothesis tests (AU, WKH, and WSH) in comparison with partition scheme and GHOST models for most datasets (Table 3), which demonstrated that the topology created by the PMSF model might be made for revealing the complicated relationships of lepidopteran superfamilies. In addition, to examine the performance of Insecta and lepidoptera datasets in tree reconstruction, we selected two topologies BUSCO30_test and lep_busco created using PMSF model with same taxon sampling for topologies test, notable exceptions to lepidoptera dataset with largest likelihood value (−22242152.35 *vs* −22263203.96) and high *p*-values (AU: 1 *vs* 0.000169; WKH: 1 *vs* 0; and WSH: 1 *vs* 0) is more suitable for delivering the high-level phylogenetic relationships discussion in context.

**Table 3.** Comparison of tree topologies created on three models using IQtree based on seven datasets.

| **Matrix** | **Models** | **logL** | **deltaL** | **bp-RELL** | **p-WKH** | **p-WSH** | **p-AU** |
| --- | --- | --- | --- | --- | --- | --- | --- |
| BUSCO30 | partition | -4963452.941 | 46.727 | 0.237* | 0.237* | 0.412* | 0.223* |
|  | GHOST | -4965014.682 | 1608.5 | 0 | 0 | 0 | 1.00e-02 |
|  | PMSF | -4963406.214 | 0 | 0.763* | 0.763* | 0.911* | 0.777* |
| BUSCO60 | partition | -12740719.55 | 540.49 | 0.0007 | 0.0005 | 0.0014 | 0.000468 |
|  | GHOST | -12742571.37 | 2392.3 | 0 | 0 | 0 | 2.60e-42 |
|  | PMSF | -12740179.05 | 0 | 0.999* | 1* | 1* | 1* |
| BUSCO70 | partition | -15609754.96 | 292.07 | 0.0005 | 0.0005 | 0.0009 | 0.000134 |
|  | GHOST | -15618021.77 | 8558.9 | 0 | 0 | 0 | 1.60e-62 |
|  | PMSF | -15609462.89 | 0 | 1* | 1* | 1* | 1* |
| BUSCO80 | partition | -19605798.24 | 0 | 0.499* | 0.502* | 0.761* | 0.506* |
|  | GHOST | -19607995.46 | 2197.2 | 0 | 0 | 0 | 6.67e-47 |
|  | PMSF | -19605798.24 | 0.00027949 | 0.501* | 0.498* | 0.741* | 0.494* |
| lep_busco | partition | -22242290.85 | 138.63 | 0.0027 | 0.0045 | 0.0076 | 0.0036 |
|  | GHOST | -22245769.9 | 3617.7 | 0 | 0 | 0 | 5.34e-63 |
|  | PMSF | -22242152.22 | 0 | 0.997* | 0.996* | 1* | 0.996* |
| noditrysia_busco | partition | -4286519.598 | 0 | 0.249* | 0.5* | 0.932* | 0.476* |
|  | GHOST | -4286519.598 | 5.48e-05 | 0.337* | 0.5* | 0.514* | 0.51* |
|  | PMSF | -4286519.599 | 0.0002471 | 0.414* | 0.496* | 0.547* | 0.476* |
| ditrysia_busco | partition | -27168915.3 | 261.14 | 0 | 0.0001 | 0.0002 | 0.00379 |
|  | GHOST | -27168976.95 | 322.79 | 0.0245 | 0.0268 | 0.0358 | 0.0236 |
|  | PMSF | -27168654.17 | 0 | 0.976* | 0.973* | 0.997* | 0.977* |

### Effect of diverse taxon sampling and none outgroups for tree reconstruction

In the present study, we retained the taxon of BUSCO completeness over 30,60,70 and 80 against insecta reference gene sets (n=1367), making four strategies to assess the effect of diverse taxon sampling for tree reconstruction in lepidoptera. The positions of 11 superfamilies (Carposinoidea, Epermenioidea, Pterophoroidea, Thyridoidea, Gelechioidea, Papilionoidea, Tortricoidea, Cossoidea, Zygaenoidea, Immoidea and Urodoidea) of ditrysia and 6 superfamilies (Palaephatoidea, Adeloidea, Nepticuloidea, Hepialoidea, Eriocranioidea and Micropterigoidea) of ditrysia that existed commonly in all trial datasets, were as the test standards for measurement. As mentioned above, the estimated phylogenetic relationships within 6 nonditrysian superfamilies were clear and stable, there was no apparent conflict along with taxon sampling (Fig. 2, Additional file 2: Figure S31). Nonetheless, for 11 ditrysian superfamilies, the positions with considerably conflicts were pretty apparent among four datasets, even though the phylogenetic signal of the datasets continued to increase, indicating that taxon composition might be a significant factor for tree robustness in phylogenetic reconstruction of ditrysia.

Furthermore, we also supplemented three lepidopteran datasets (lep_busco, ditrysia_busco and nonditrysia_busco) with none Trichoptera taxa to test whether outgroups affected reconstructions of deep nodes in the ingroups. For so-called nonditrysia, relationships along the backbone nodes and between its superfamilies were quite consistent in the concatenation and MSC analyses, with the nearly same topology not only found in same insecta (BUSCO30 *vs* BUSCO30_test) or lepidopteran (lep_busco *vs* nonditrysia_busco) datasets, but also appeared among different datasets (Additional file 2: Figure S1-S3, S13-S15, S19-S22, S25, S29). However, for species-rich ditrysia, there was lacking of consensus stable topologies within its superfamilies (BUSCO30 *vs* BUSCO30_test), while corresponding measures (same dataset and phylogenetic model) were done (Additional file 2: Figure S3, S22). What reason is causing this significant discrepancy for ditrysian phylogeny? Is it controlled by more distant outgroups and abundant characters? To answer this question, we did a test using two lep_busco and ditrysia_busco datasets, one contained almost all taxa within key lepidopteran clades, and the other contained only ditrysian taxa but the site was more than 400 thousand (Table 2). As expected, it was clearly showed that inconsistent topologies were still remained (Additional file 2: Figure S13-S18, S29-S30), with no effect by distant lineages and data quality. Taken all together, the results strongly indicated that the outgroups could result in biased ingroup topologies on phylogenetic reconstruction of ditrysia to a certain extent, whereas have no effects on that of stable nonditrysia.

### Discordance among gene trees

To more systematically for quantifying genealogical concordance of coalescent and ASTRAL trees, we implemented the gene concordance (gCF) and site concordance (sCF) factor compared gene tree and species tree topologies to determine the number of gene or sites that conflict at each node. Almost all datasets yielded highly ultrafast bootstrap and maximum LPP values for most branches in coalescent and ASTRAL analyses (Additional file 1: Table S6-S7, Additinal file 2: Figure S3, S6, S9-S10, S15, S18, S25-S30). However, it was clearly showed that high bootstrap and LPP did not always coincide with high gCFs and sCFs. The gCF values (39.95-44.07) were much lower than the sCF values (51.42-52.56) in average among the trial datasets (Additional file 1: Table S7), which showed that the sites that support this topology are more prominent than the genes. Moreover, it was found that the branches in the backbone were shorter branches in the tree and short branches had the low CF values (Additional file 2: Figure S3, S6, S9-S10, S15, S18, S25-S30). Of note, the CF values had the significant divergence across different taxonomies of lepidoptera (Fig. 3, Additional file 2: Figure S32). For example, the gCFs and sCFs values in genus-level were ranged from of 48.45-52.98 and 55.61-57.39 in average, whereas that in family-level were 13.81-18.78 and 36.84-39.32 in average, and in superfamily-level were 3.88-12.05 and 35.14-37.52 in average (Additional file 1: Table S7), which indicated that high CF values in accord with high bootstraps might be more remarkable in shallower taxonomic level of lepidoptera.

**Figure 3.**
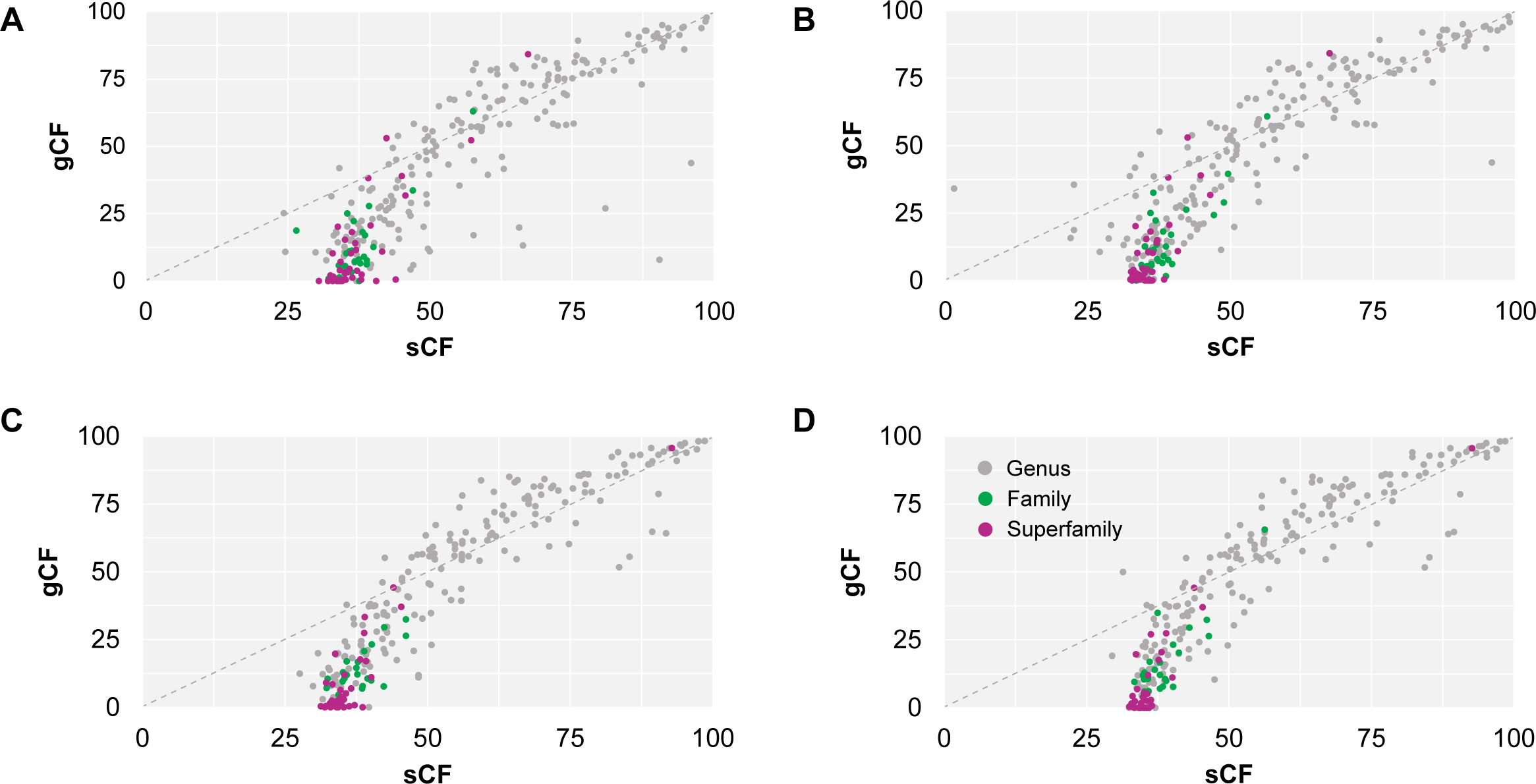
The links of gene concordant (gCF) and site concordant (sCF) factors across all nodes of the ML and ASTRAL trees. **(A)** Concatenated ML tree under the PMSF model of BUSCO30 dataset. **(B)** ASTRAL tree of BUSCO30. **(C)** Concatenated ML tree under the PMSF model of lep_busco dataset. **(D)** ASTRAL tree of lep_busco. The colors in the figure represent three different classification levels (superfamily, family and genus), respectively.

### Phylogenetic relationships of lepidopteran superfamilies

In this study, phylogenetic reconstruction was performed using maximum-likelihood (ML) and Bayesian inference (BI) methods based on eight amino-acid datasets. ML analysis completed first, while BI analysis with large memory demand (RAM usage for lep_busco dataset was more than 790 G) in PhyloBayes has been still running more than two months, without convergence, so that we have to terminate program due to the heavy computational burden. The results showed that multiple tree topologies were generated by using a diverse set of analytical methods (Additional file 2: Figure S1-S30). The structures of BI trees with strong supports were nearly identical with that of the ML trees (PMSF) of corresponding datasets (Additional file 2: Figure S3, S15, S23, S24), whereas the ASTRAL topologies were weakness for the reason that the shorter genes within datasets could not recover the accurate relationships of lineages with divergences more than a hundred million years ago (Table 2). As for the rest of the topologies, various conflicting structures still hindered our understanding the complicated phylogenetic relationships of lepidoptera. We carried out topology test for our better decision which topology with good performance was the suitable for delivering the discussion in context. It was clear that the lepidoptera datasets with the PMSF model in ML analysis exhibited better robustness for tree topologies than insecta datasets (Table 3). All things considered, the tree topology of lep_busco dataset based on the PMSF model in ML analysis was hereafter adopted for discussion the phylogenetic relationships of lepidopteran superfamilies (Fig. 4 and Fig. 5). According to the tree, 10 key clades (Eight marked in Fig. 3 and Fig. 4, and two marked in Fig. 2A) span the whole breadth of the lepidopteran tree of life were all present in resulting topologies with high branch supports (UFB = 100, SH = 100). The backbone of basal lepidopteran superfamilies was stable, but that of middle-upper lepidopteran superfamilies had some considerably divergence in comparison with the previous studies (Fig. 1, Fig. 4).

**Figure 4.**
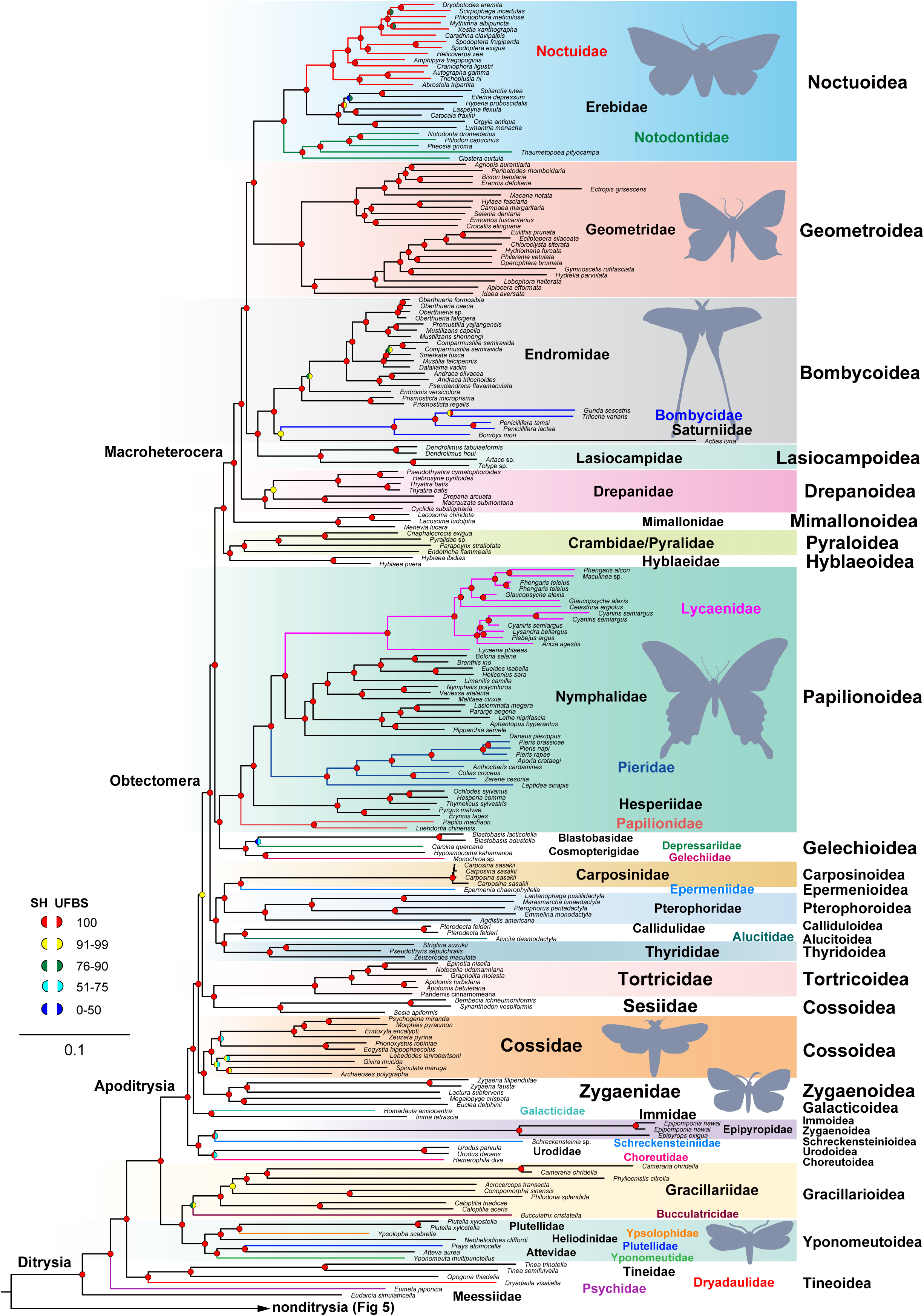
Phylogenetic tree of superfamily relationships within ditrysian Lepidoptera inferred from maximum likelihood of the lep_busco dataset based on PMSF model in IQtree using 1,017 universal single copy orthologues for 250 taxa. This is a partitioned tree divided from lep_busco tree, simplified to present the major lineages of ditrysia. A complete tree for all Lepidoptera is connected with Figure 5 and is shown in Additional file 2: Figure S15. Colored labels at the nodes represent the summarized status of bootstrap (SH-aLRT and UFBS) values in lep_busco tree.

**Figure 5.**
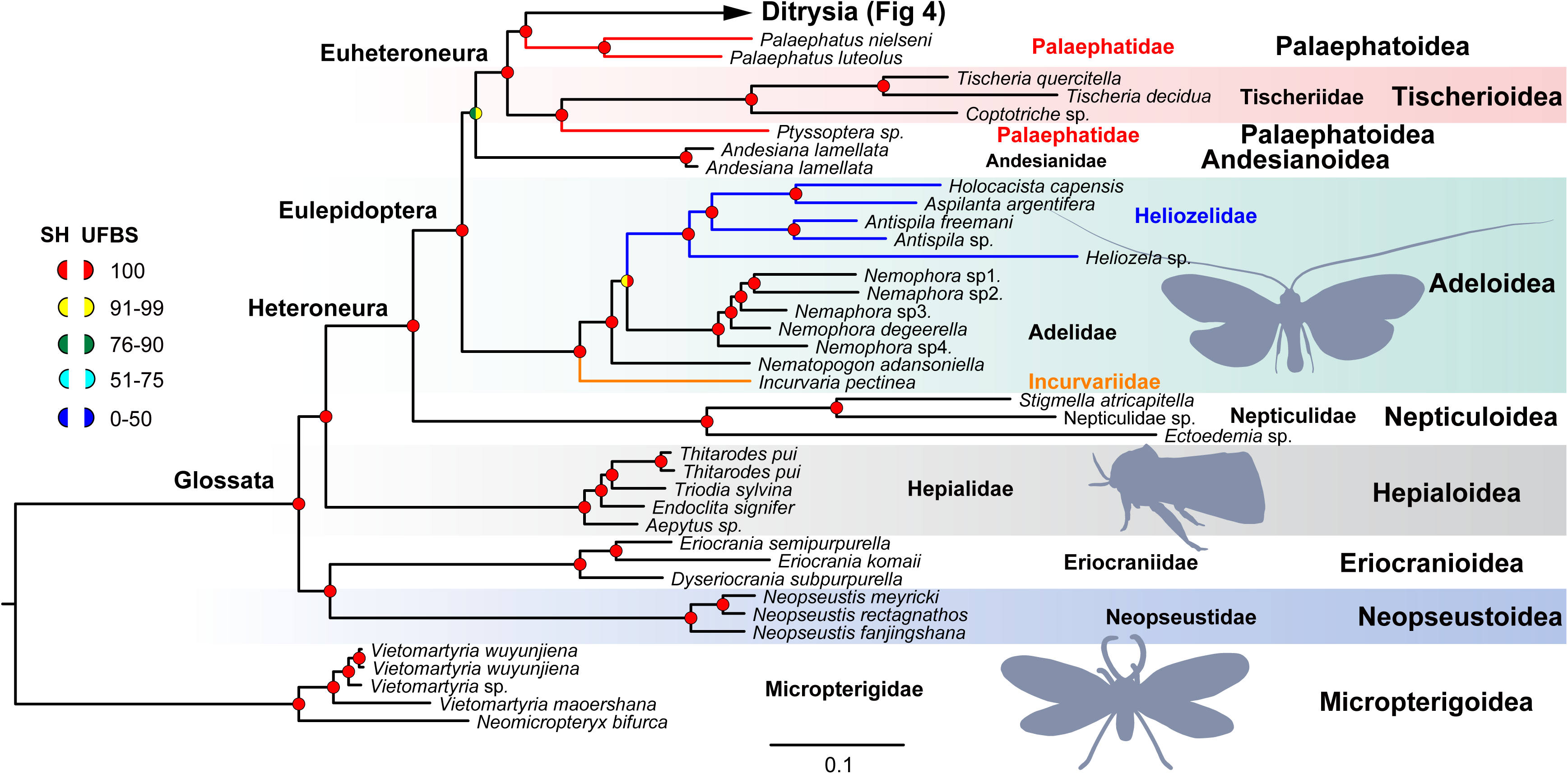
Phylogenetic tree of superfamily relationships within nonditrysian lepidoptera inferred from maximum likelihood of the lep_busco dataset based on PMSF model in IQtree using 1,017 universal single copy orthologues for 250 taxa. This is a partitioned tree divided from lep_busco tree, simplified to present the major lineages of nonditrysia. A complete tree for all Lepidoptera is connected with Figure 4 and is shown in Additional file 2: Figure S15. Definitions of labels on each node are the same as in Figure 4.

Within ditrysia (Fig. 4, Additional file 2: Figure S15), the non-monophyletic Tineoidea were followed by a clade consisting of Gracillarioidea and Yponomeutoidea in the base. Choreutoidea and Urodoidea was as the sister group to Schreckensteinioidea and Zygaenoidea part (Epipyropidae), all of them assemblage formed the basal branches of Apoditrysia with strong support (UFB = 100, SH = 100). Moreover, Zygaenoidea was not monophyletic, and the remainder of the clade being sister to Cossoidea (Cossidae). Cossoidea was also particularly intriguing as it displayed different positions, the other (Sesiidae) was sister to Tortricoidea in the upper of Apoditrysia. Thyridoidea + Alucitoidea + Calliduloidea was formed an association, which was placed as a sister-group relationship with another association of Pterophoroidea + Epermenioidea + Carposinoidea in Obtectomera clade (UFB = 100, SH = 100). It was worth noting that Papilionoidea was recovered as sister group to Gelechioidea with strong support (UFB = 100, SH = 100). For the upper topologies of lepidoptera, the relationships among each superfamily were relatively stable. Pyraloidea was as the sister group relationship with Hyblaeoidea outside of the Macroheterocera. Noctuoidea + Geometroidea was recovered in the place of Bombycoidea + Lasiocampoidea with high support values (UFB = 100, SH = 100). Within so-call nonditrysia (Fig. 5, Additional file 2: Figure S15), Micropterigoidea was placed as the base to all other Lepidoptera. Neopseustoidea + Eriocranioidea was subsequently placed sister group to the remaining Lepidoptera. For Eulepidoptera, Adeloidea was formed an independent clade outside of Andesianoidea. Palaephatoidea was split into two clades, one clade (*Ptyssoptera*) was recovered as sister group of Tischerioidea and the other (*Palaephatus*) was sister to the ditrysia with the highest statistical support (UFB = 100, SH = 100).

## Discussion

### Data effectiveness and tree topology robustness

Benchmarking universal single-copy orthologs (BUSCOs) are a set of conservative benchmarking genes of universal single-copy orthologs identified using OrthoDB [25], which are commonly used for quantitative assessment of genome assembly and annotation completeness based on evolutionarily informed expectations of gene content [26, 27]. The potential of BUSCOs is further explored to downstream phylogenetic inference [26], whose effectiveness with high branch supports is tested and verified not only in diverse high-level lineages, such as insects [50], yeasts [51] and spiders [52], Entomobryomorpha [30], and Hemiptera [53]; but also in some low-level lineages Halictini [54], Membracoidea [55] and Chrysopidae [29]. In this study, 8 coalescent datasets with higher parsimony information sites yielded 24 phylogenetic trees with high branch supports (Table 2, Additional file 2: Figure S1-S24). All of these phylogenetic reconstructions were clearly shown that the relationships of nonditrysian superfamilies were almost unambiguous each other (Fig. 2, Additional file 1: Table S6, Additional file 2: Figure S1-S24), and the similar results were also observed in previous studies (Fig. 1), which strongly indicated the relationships among the paraphyletic assemblage of so-called nonditrysian lepidoptera are stable, and our datasets either insecta or lepidoptera are effective for reconstructing the tree of life of lepidoptera.

Evolutionary models play an essential role in systematic biology, which could be simplifications of the evolutionary processes for facilitating biologists better understand the origin and evolution of lineages [56–58]. Here, we carried out phylogenetic inference using a diverse set of models (partitioned maximum likelihood, heterotachy model, site heterogeneous model, and multispecies coalescent model) reduced systematic biases to reveal the relationships among lepidopteran superfamilies (Additional file 2: Figure S1-S30), and topologies tests were done for quantification the performance of three ML-based models. The results showed a variety of inconsistent tree topologies among ditrysian superfamilies and relatively stable backbone within nonditrysian superfamilies (Fig. 2, Additional file 1: Table S6, Additional file 2: Figure S1-S30). PMSF model got a good performance with higher *p*-values in compared with partitioning and GHOST model in almost all hypothesis tests (Table 3). Recent model evaluation demonstrated that site-wise heterogeneity effects leaded to a serious long-branch attraction (LBA) bias are extreme than protein-wise heterotachy effects in phylogenomic analyses [59]. Therefore, we concluded that site heterogeneity might be an important source of bias given rise to topologies divergence of ditrysia in phylogenomic reconstruction of lepidoptera. Furthermore, incomplete lineage sorting (ILS) is also another significant bias providing misleading information for phylogeny on account of topologies of gene tree inconsistent with that of species tree [60–62]. Differences observed in the present study between the ASTRAL and concatenated trees do not necessarily mean that there is gene tree versus species tree conflict (Additional file 1: Table S6, Additional file 2: Figure S1-S30). One possible reason is that the shorter gene fragments (average length is less than 340 pb) within these datasets could not be able to recover the accurate relationships of lineages with divergences more than a hundred million years ago (Apoditrysia are estimated to be about 130 Ma [35]), and another might be that limited number of loci (number of loci are less than 1266) cannot afford plenty robust statistics. Therefore, to some extent, our ASTRAL trees are weak and could not deliver phylogenomic needs in deep-level relationships of lepidoptera.

Concordance factors (CF) are significant criterions for quantifying genealogical concordance in phylogenomic datasets to gain a deeper understanding of how well different genes support the different topologies [63]. For most branches in present tree topologies, the gCF values were much lower than the sCF values (Additional file 2: Figure S3, S6, S9-S10, S15, S18, S25-S30), suggesting that the sites that support the topology are scattered across the different genes. Additionally, as has been demonstrated in higher-level phylogenetic inference of other lineages, such as treefrogs [64], Membracoidea [55], and spiders [65], concordance factors in many branches with perfect supports have quite low CF value. These similar results not only observed in our study (Additional file 1: Table S7), but also appeared in other lepidopteran phylogenomics [49]. It is worth noting that the CF values had the significant divergence in different taxonomic levels of lepidoptera (Fig. 3, Additional file 2: Figure S32), the CFs values in genus-level were tend to higher than that in family-level or superfamily-level. As a result, all the above findings strongly indicated that adopting the concordance factors for farther assessing and supplementing branch supports might be ineffective in higher-level phylogenetic reconstructions, and this phenomenon may be more noticeably with the taxon increased.

### Effect of taxon sampling for tree reconstruction

Taxon sampling has a positive effect on tree topology robustness in phylogenetic reconstruction [66, 67]. Appropriate and extensive taxon sampling is one of the most important determinants of accuracy of inferences about evolutionary processes [68]. Although a large number of taxa create a more complex computational problem for phylogenetic analysis, its excellent performance in reducing the effect of LBA has been verified and accepted by a large number of phylogenetic estimations [69–74]. Furthermore, it is a widely recognized that increased character data can enhance phylogenetic signal getting a good resolution and support and reduce stochastic error in large-scale phylogenomics [75]. In this study, we considered both the sampling of taxa, as well as the number of characters, in designing a series of datasets for phylogenomic studies. The positions of 11 ditrysian and 6 nonditrysian superfamilies commonly existed in all trial datasets were as the test standards for measurement. It was obvious showed that the relationships among 6 nonditrysian superfamilies were unambiguous, whereas among 11 ditrysian superfamilies remained equivocal even with increasing the total number of characters in datasets (Fig. 2, Additional file 2: Figure S31). Previous studies indicated that the discordant placement of the superfamily Gelechioidea was caused by the poorly taxon sampling [49]. However, this effect does not only occur in Gelechioidea, our results showed that the discordant phylogenetic positions of almost all ditrysian groups, especially for apoditrysian groups, have been infected by this insufficient sampled. Although increasing the total number of characters could enhance the phylogenetic accuracy in previous discussions [76–78], this homoplasy is not clear enough in our results. For example, we employed a series of analytical methods reducing system errors to dissect our datasets and demonstrate that there is sufficient phylogenetic signal to resolve nonditrysian phylogeny, but not all ditrysian phylogeny, even with increasing numbers of sites and exceeding 400,000, which demonstrated that taxon sampling is the mainly factor contributing on topological accuracy of ditrysian phylogenetic reconstruction. Moreover, previous assessment has shown that using fewer genes could mitigate the effect of long-branch attraction and get acceptable support for the correct topologies on a more densely sampled yeast phylogeny when taxon sampling is increased [79]. Therefore, strongly advise for researchers to increase taxon sampling of lepidopteran group in the future phylogenetic inference, especially for Ditrysia, the most species rich lepidopteran clade containing 98% of the extant species. If the density of taxon sampling is increased, additional internal nodes could reveal undetected substitutions and improve estimates of branch lengths [68], perhaps a certain amount of data could show a clearer and more accurate phylogenetic estimation of Ditrysia, which would help settle the arguments of the placements among its superfamilies.

### Phylogenetic relationships of lepidopteran superfamilies

The lepidopteran tree of life has been controversial for a long time (Fig. 1), with a number of conflicts between different studies and even different analyses of the same data [35, 46, 49]. The most unresolved part of the Lepidoptera tree of life is Ditrysia [80]. Within Ditrysia, the superfamily Tineoidea was recovered repeatedly with three separate lineages branching off sequentially to the rest of Ditrysia in this study, first genus *Eudarcia* (Meessiidae), then genus *Eumeta* (Psychidae) and finally the families Dryadaulidae + Tineidae together recovered as sister to all other Ditrysia (Fig. 4). However, in our ditrysia dataset, Tineoidea was recovered as a monophyletic group with strong supports (Additional file 2: Figure S16-S18, S30), similar result was also observed in Heikkilä et al. (2015) phylogeny that combined the eight genes (COI and seven nuclear genes) with morphological characters [40]. Previous phylogenetic inferences based on much better taxon sampling with fewer genes (19 nuclear protein-coding genes) across Tineoidea or large-scale phylogenomics with one consent showed that the Tineoidea were paraphyletic with multiple clades and the family Tineidae being sister to all other Ditrysia [35, 41, 48, 49]. Thus, all of these studies clearly indicated that the relationships within the superfamily Tineoidea are questionable and need to farther increase taxon sampling getting a robust and stable phylogenetic framework for redefinition to render the superfamily monophyletic.

Within the rest of Ditrysia, the position of Yponomeutoidea + Gracillarioidea as sister to Apoditrysia is well supported in all of the studies (Fig. 4) [35, 39, 47–49]. There is a disagreement across our datasets with previous phylogenetic inferences in the position of Choreutoidea, Urodoidea, Schreckensteinioidea and Zygaenoidea (Epipyropidae) outside of Obtectomera as their placement to some extent depends on the datasets as well as analysis type. Choreutidae is monotypic family belong to Choreutoidea of Apoditrysia described by Minet (1991) [81, 82], whereas molecular phylogeny based on gene marker showed that it is polyphyletic and had been placed in different superfamilies [38, 39, 83]. In this study, Choreutoidea and Urodoidea is formed a sister group in line with Kawahara et al. (2019), Mayer et al. (2021) and Rota et al. (2022) [35, 48, 49], which was not shown the polyphyletic clades of Choreutoidea due to the poor sampled. Schreckensteinioidea and Zygaenoidea (Epipyropidae) recovered as sister group is the first observed here. In previous studies, Schreckensteinioidea is usually sister to Urodoidea [40], and Zygaenoidea is monophyletic as the immediate sister to Cossoidea shown in plenty of literatures [35, 40, 46–49], whereas it was split into two clades in this study – one (Epipyropidae) was closely related to Schreckensteinioidea and another (Zygaenidae) was sister to Cossoidea (Cossidae). Equally confusing is that Cossoidea stand out as being consistently recovered as two separate lineages. In addition to the above mentioned Cossidae, the other (Sesiidae, belonged to the superfamily Sesioidea in the past) was sister to Tortricoidea. Although the studies of fewer genes with much better taxon sampling (weak supports) [40] and genome-scale phylogenies (strong supports) with poor sampled [35, 47–49] are consistently demonstrated a monophyletic group and closed relationships of Cossoidea and Zygaenoidea each other, a disagreement (strong supports) was shown here using different analyses under relatively more genomic data. Therefore, for these two groups, there is an urgent need for farther increasing taxon sampling and adopting large-scale genomic data to clarify clearly their monophyletic in the future phylogeny. Within Obtectomera, the case of superfamily Pterophoroidea is great interesting, as its position has changed wildly in previous studies. Mutanen et al. (2010) found it forms an assemblage contained with Alucitoidea, Epermenioidea, Schreckensteinioidea, Copromorphoidea and Urodoidea outside of Obtectomera [38]. Regier et al. (2013), on the contrary, placed them (along with Carposinoidea) as sister to Papilionoidea within Obtectomera [39]. Similarly, in the phylogenomic studies, our results are similar to Bazinet et al. (2013), Kawahara et al. (2019) and Mayer et al. (2021) showed that Pterophoroidea was formed an association with multiple superfamilies (Thyridoidea, Alucitoidea, Calliduloidea and so on) within Obtectomera [35, 45, 48], whereas Kawahara et al. (2014) and Rota et al. (2022) found Pterophoroidea to be sister to Urodoidea or Carposinoidea outside of Obtectomera [46, 49]. Moreover, the others controversial parts of Obtectomera tree of life are Gelechioidea and Papilionoidea. Gelechioidea was sister to Papilionoidea within Obtectomera in the present study (Fig. 4). However, previous studies found it forms an alone clade [39, 47–49] or association with some superfamilies [35, 40, 45] (Thyridoidea, Alucitoidea, Pterophoroidea and so on) within Obtectomera, whereas Mutanen et al. (2010) showed that it is outside of the association of Thyridoidea + Calliduloidea + Papilionoidea + Hesperioidea + Hedyloidea within Apoditrysia [38]. Papilionoidea (butterflies), as phytophagous insects with successful evolutionarily, play a central role in the study of speciation, community ecology, biogeography, climate change, and plant-insect interactions [32, 84]. Recent research based on dense taxon sampling (nearly 2,000 species representing 92% of all butterfly genera) aggregated global distribution records and larval host datasets presents a comprehensive phylogenetic framework of butterflies up to now [85], but the position of that within Lepidoptera are still unresolve. Some phylogenetic inferences showed that it forms an assemblage with a few superfamilies (Pterophoridae, Calliduloidea, and Thyridoidea) outside of Pyraloidea + Hyblaeoidea [38-40, 45, 47, 49]; and others showed an independent clade that closed to Zygaenoidea + Cossoidea away from of Pyraloidea + Hyblaeoidea [35, 47]. Clearly, for Obtectomera group, a more careful investigation is needed to establish to put an end to the debate over the positions of confusion among its superfamilies. Within Macroheterocera, disagreement also exists in the placement of the groups Noctuoidea, Geometroidea, Bombycoidea and Lasiocampoidea. Noctuoidea is sister to (Bombycoidea + Lasiocampoidea) showed in Regier et al. (2013), Heikkilä et al. (2015) and Rota et al. (2022) [39, 40, 49], while Geometroidea replaced of Noctuoidea is sister to (Bombycoidea + Lasiocampoidea) by Kawahara et al. (2019) and Mayer et al. (2021) [35, 48]. In this study, Geometroidea + Noctuoidea was as sister group to Bombycoidea + Lasiocampoidea in line with Kawahara et al. (2014), Breinholt et al. (2018) and Kim et al. (2020) [44, 46, 47].

Relationships among the paraphyletic assemblage of so-called nonditrysian lepidoptera appear to be settling down based on plentiful studies with more taxa and genes [35, 39, 42, 46, 48, 49, 86]. It is clearly now that Micropterigoidea are sister to the rest of Lepidoptera, followed by Agathiphagidae and then Heterobathmiidae [49, 86]. Previous evaluations with a handful of genes suggested that Agathiphagoidea were sister to Heterobathmioidea [38, 42, 46], but this result is strongly incongruent with morphological data [82]. In this study, there is no ability to check this arrangement on reason that Agathiphagoidea samples was not used due to the weak completeness (Additional file 1: Table S4). Moreover, the position of other small superfamilies, Neopseustoidea, Eriocranioidea, and Hepialoidea have been elusive for a long time. Neopseustoidea and Eriocranioidea form a sister group outside of Hepialoidea were found in the present study with strong supports (Fig. 5), which are in line with Kawahara et al. (2019) and Rota et al. (2022) [35, 49], whereas they are clustered an assemblage with Lophocoronoidea in Mayer et al. (2021) (see Fig. 1) [48]. For the rest nonditrysian superfamilies, Adeloidea and Andesianoidea are recovered as sister group in Kawahara et al. (2019) and Mayer et al. (2021) [35, 48], whereas they form an independent clade each other in Rota et al. (2022) and this study. Palaephatoidea, a group with Gondwanan distribution, stand out as being consistently recovered as two separate clades [49]. For this lineage, an important question of which group is the sister to Ditrysia is still controversial [42, 49]. In previous study the Palaephatoidea genera *Azaleodes*, *Metaphatus* and *Ptyssoptera* are closed related to Tischeriidae, which are together sister to Ditrysia whereas the genus *Palaephatus* forms a separate lineage that is sister to all nonditrysian groups [49]. However, this relationship is reversed and *Palaephatus* is recovered as the immediate sister to Ditrysia (Fig. 5), is shown in the present study, Kawahara et al. (2019) and Mayer et al. (2021) [35, 48].

In summary, our phylogenomic trees contain the largest sampling of superfamily (37 superfamilies) of lepidoptera to date, and present the good results of ditrysian and nonditrysian superfamilies with high branch supports (Additional file 1: Figure S1-S30), as well as get some surprising discovers in comparison of previous studies [35, 38–42, 45, 46, 48, 49, 80, 86]. However, as discussion before, taxon sampling is an important factor in tree robustness of ditrysian phylogenetic reconstruction. 37 of 43 superfamilies were sampled but 6 superfamilies Agathiphagoidea, Acanthopteroctetoidea, Lophocoronoidea, Mnesarchaeoidea, Simaethistoidea, and Whalleyanoidea were not sampled in the present phylogeny. Thus, our phylogenetic tree could only represent the current understanding of the lepidopteran tree of life. The relationships within the species-rich and relatively rapid radiation Ditrysia and especially Apoditrysia remain unresolved and elusive for some time. With the continuous advancement of Earth BioGenome Project (EBP), more and more genomic resource of Lepidoptera will be available in public database. How to obtain sufficient phylogenetic information from massive data may still be a problem that researchers need to consider carefully. Current phylogenomics is now generally accepted as the use of more and more genes in large-scale level for precise phylogenetic reconstruction [4, 75, 87]. However, a recent study showed that lepidopteran insects have the high average number of horizontal gene transfer (HGT) acquired genes [88], which make us think seriously about whether the genes with high phylogenetic signal and bootstraps must be suitable for delivering phylogenetic needs, and could they also be noise in the actual phylogeny of lepidoptera? Thus, our hope is that significantly increasing taxon sampling across the ditrysian lineages while adopting lineage-specific genes for phylogenomic reconstruction would lead to further resolution in the lepidopteran tree of life.

## Conclusions

In the present study, we increased taxon sampling among Lepidoptera (40 superfamilies and 76 families contained 286 taxa) and filtered the unqualified samples, then acquired a series of large amino-acid datasets derived from genomes and transcriptomes against the insecta (n=1,367) and lepidoptera (n=5,286) reference gene sets for phylogenomic reconstructions. Using these datasets, we explored the effect of different taxon sampling for tree topology over considering a series of systematic errors using ML and BI methods. Moreover, we also tested the effectiveness in topology robustness among the three ML-based models. The results showed that taxon sampling is an important factor in tree robustness of accurate lepidopteran phylogenetic estimation. Site-wise heterogeneity leading to a serious long-branch attraction (LBA) is a significant source of bias given rise to topologies divergence of ditrysia in phylogenomic reconstruction of lepidoptera. Moreover, our phylogenetic tree is by far the most comprehensive to reveal the relationships among lepidopteran superfamilies, but limited by taxon sampling, it could only represent the current understanding of the lepidopteran tree of life. The relationships within the species-rich and relatively rapid radiation Ditrysia and especially Apoditrysia remain poorly unresolved and need to further research.

## Methods

### Taxon sampling and DNA extraction

A total of 286 taxa representing 40 of 43 currently recognized lepidopteran superfamilies and 76 of 137 families was used in the present study. Among them, 52 individuals representing 48 species (12 superfamilies,17 families) were selected from specimens cataloged in Insect Museum of Hunan Agricultural University used for newly sequenced, and the detail collection information were showed in Additional file 1: Table S1. For others, genomes assembled at the chromosome or scaffold level of 111 taxa representing 14 superfamilies (27 families) were downloaded from the genome database of NCBI (https://www.ncbi.nlm.nih.gov/genome) (Additional file 1: Table S2), and the remain taxa (119 taxa) representing 32 superfamilies (47 families) were from the SRA database of NCBI (https://www.ncbi.nlm.nih.gov/sra/) for farther genome or transcriptome assembly (Additional file 1: Table S3, Table S4). Furthermore, five genomes of Trichoptera were also from NCBI using as outgroups for phylogenomic analysis in this study. All taxa from NCBI were rechecked and adjusted their current locations in lepidoptera according to the latest taxonomy by literature review.

For those newly sequenced individuals, all specimens were identified on the basis of morphological characteristics via external characters and male genitalia, then checked and confirmed its identifications by cytochrome c oxidase subunit 1 (COI) DNA barcodes which matched in NCBI and BOLD databases at least family level. Total genomic DNA was extracted from the leg tissue of a single specimen (total body of larvae) following the protocol provided by SteadyPure Universal Genomic DNA Extraction Kit Ver.1.0 (Changsha, Hunan, China), with some modifications: incubation time was increased to 24h with extra addition of 10 μl of proteinase K. DNA was preserved at −20 ℃ and used for sequencing.

### Genome sequencing and assembly

Libraries of an insert size of 350 bp were constructed and sequenced (2 × 150 bp) on the Illumina Novaseq 6000 platform at Berry Genomics (Beijing, China). For each library, 6 Gb of clean data were obtained for each individual, which were usually sufficient to generate enough number of targeted loci for phylogenomic inference. Raw data was assessed the quality of the generated using FastQC v0.11.9 (https://www.bioinformatics.babraham.ac.uk/projects/fastqc/), then filtered using fastp v0.23.2 [88], with adapter sequences trimmed referring to the self-provided Illumina adapter sequence database, also leading and trailing bases with quality below 30 were removed for each read. Subsequently, Trinity v2.2.1 [89] was used to perform *de novo* assemblies for transcriptome data with default settings, and redundant contigs from each sample were reduced using CD-HIT v4.8.1 [90] with the parameter ‘-c 0.9’. *de novo* genome assembly was performed using Spades v3.15.3 [91] with kmer values of 21,41,61,81,101 and 121, and gap filling for each draft genome was performed using GapCloser v1.12 [92]. All draft genomes assembled were filtered the contigs less than 1000 bp in length using BBmaps v38.93 (https://sourceforge.net/projects/bbmap/) for fast and accurate gene prediction in following analysis. The completeness of each assembly (genomes or transcriptomes) was assessed using BUSCO v.5.2.2 [27] with the parameter ‘-m geno’, calling the augustus [93] module for gene prediction using the Insecta (n=1368) lineage set.

### Dataset generation and Sequence alignment

Benchmarking universal single-copy orthologues (BUSCOs) were extracted from genomes or transcriptomes with BUSCO v5.5.2 [27] against the Insecta (n=1368) and lepidoptera (n=5286) reference gene set. However, for some lineages (e.g. Agathiphagidae and Lophocoronoidea), the completeness of each assembly was less than 30% against the Insecta (n=1367) lineage set, so that there were not enough universal single-copy orthologues (USCOs) for phylogenetic inference in the present study. We excluded those taxa with completeness below 30% (∼ 418 genes) to ensure adequate phylogenetic signal. Moreover, for the species *Whalleyana vroni* (SRA number: SRR11799523) belong to superfamily Whalleyanoidea, we also excluded this sample on account of failed genome assembly in our server (assembly it required more than 1.5T memory). Finally, 263 taxa representing 37 of 43 currently recognized lepidopteran superfamilies and 67 of 137 families were selected and adopted for following phylogenetic inference.

To reduce compositional heterogeneity, BUSCOs amino acid was used for subsequent analyses. We generated five insecta matrices (BUSCO30, BUSCO30_test, BUSCO60, BUSCO70, BUSCO80) and three lepidopteran matrices (lep_busco, ditrysia_busco and nonditrysia_busco) to assess the effect of diverse taxon sampling for tree reconstruction in lepidoptera (Table 2). For genes selecting, the completeness of genome or transcriptome was various due to the different depth and coverage of sequencing in each taxon, and the composition of genes was different even within the same completeness. Thus, the gene that occurred or existed in more than 90% of all taxa was defined as common gene used for matrix generation. Protein sequences of each locus were aligned by using MAFFT v7.505 [94] with the L-INS-I algorithm, and were trimmed by using BMGE v1.12 [95] with the stringent parameters ‘-m BLOSUM90 -h 0.4’. Trimmed alignments were concatenated by using FASConCAT-g v1.05.1 [96]. We further excluded those alignments that violated the SRH (stationary, reversible and homogeneous) assumptions using ‘--symtest’ implemented in IQ-TREE v2.2.0.3 [97, 98]. Selection of genes with strong phylogenetic signal, which could be reflected by high parsimony informative sites (PISs) and bootstrap supports, could reduce topological incongruence between gene trees and species trees and improve resolution particularly for deep nodes [99–101]. The PISs of all locus were performed by PhyKIT v1.11.10 [102], then we retained the loci that the percentage of PISs greater than 50. Subsequently, individual gene trees of these loci were inferred by using IQ-TREE with the ‘EX_EHO’ mixture protein model and 1,000 ultrafast bootstraps (UFBoot2) [103], and we selected BUSCOs of average UFBoot2 values greater than 50 to generate the matrices. The basic statistics for the captured BUSCOs of the eight matrices were shown in Table 2.

### Phylogenetic reconstruction and Topology robustness

Phylogenomic analyses was performed using maximum-likelihood (ML) and Bayesian inference (BI) methods initially on four Insecta amino-acid datasets of selected universal single-copy orthologous proteins. However, we could not be sure that the BUSCO of Insecta lineage set was suitable for revealing phylogenetic relationships of lepidoptera due to the distant lineages with more conservative universal single-copy orthologous, especially for that of rapid radiation lineages Ditrysia [48]. We supplemented with three datasets (lep_busco, ditrysia_busco, and nonditrysia_busco) whose BUSCOs were gained against the lepidoptera reference gene set to verify the above hypothesis. Incorrect phylogenomic trees might derive from biological and methodological sources of systematic error [5, 86], such as compositional bias [104], missing data, heterotachy (i.e., rate variation across sites and lineages) [105], rate heterogeneity [106], and incomplete lineage sorting (ILS) [107, 108].

We carried out phylogenetic inference using a diverse set of analytical methods to account for these sources of bias. Measures aimed at mitigating compositional bias and missing data were implemented during matrix generation. For the BUSCO amino acid matrices, phylogenetic trees were inferred by using partitioned maximum likelihood (ML), heterotachy model, and multispecies coalescent model methods. For the partitioned ML analyses, partitioning schemes and substitution models were selected based on ModelFinder [109] implemented in IQ-TREE by employing the relaxed hierarchical clustering algorithm ‘-rclusterf 10’ [110]; estimations were restricted to a subset of amino acids models. Heterotachy ML reconstructions were performed by using the General Heterogeneous evolution On a Single Topology (GHOST) model in IQ-TREE [111]. Node supports in all ML analyses were calculated by using 1,000 Shimodaira-Hasegawa-like approximate likelihood ratio test (SH-aLRT) replicates and 1,000 UFBoot2 bootstraps [103, 112]. SH values and UFBS values were considered strong when higher than 80% and 95%, respectively. The multi-species coalescent model (MSCM) addressing ILS was run by using ASTRAL-III v5.7.8 [113, 114]; given a set of unrooted gene trees, branch supports were estimated using local posterior probabilities (LPP) on quadripartitions with default hyperparameter inputs [115]. Individual gene trees were obtained from the previous steps of matrix generation. To further assess how well different genes support the best topology, we measured gene concordant (gCF) and site concordant (sCF) factors using IQ-TREE to comprehensive measures of underlying agreements or disagreement among sites and genes for supporting topological arrangement [116].

To mitigate long-branch attraction (LBA) artefacts, we applied site-heterogeneous models in both ML and Bayesian inference contexts. The posterior mean site frequency (PMSF) [62] model was performed for each BUSCO matrix by specifying a profile mixture model with ‘LG+C20+FO+R’ for Insecta datasets and ‘LG+C10+FO+R’ for lepidoptera datasets in IQ-TREE, respectively. The corresponding partitioned ML tree was treated as an initial guide tree. The resulting PMSF tree was used as a new guide tree, leading to new PMSF profile and a new tree. The PMSF tree inference process was iteratively repeated twice, then the tree with the largest log-likelihood score was adopted as the final tree for delivering discussion in the contexts. Bayesian inference by using PhyloBayes MPI v1.8c [117] was only performed for datasets BUSCO30 and lep_busco due to the computational burden in the present study. Two separate chains were run for 3,160-3,443 generations of BUSCO30 dataset and 1,723-1,725 generations of lep_busco dataset under the CAT+GTR model [118] by using a starting tree derived from the partitioned ML analyses, respectively. The two chains converged on the same topology in all MCMC samples after removing the first 2,600 generations for BUSCO30 dataset and 1,000 generations for lep_busco dataset as the burn-in, respectively.

Phylogenetic reconstruction was performed using maximum-likelihood (ML) method based three complex models (partition, GHOST and PMSF) in IQtree. However, we could not be ensured under which models the reconstruction could get a good performance for help better our understanding the complicated relationships of the butterflies and moths due to the divergent topologies. We tested the resultant three alternative topologies with the corresponding matrix using approximately the unbiased (AU), weighted Kishino-Hasegawa (WKH), and weighted Shimodaira-Hasegawa (WSH) tests under the PMSF model ‘LG+C20+FO+R’ for Insecta datasets and ‘LG+C10+FO+R’ for lepidoptera datasets in IQ-TREE, respectively. The tree estimated from previous PMSF inference was used as the guide tree and the starting tree.

## Abbreviations

BUSCO: Benchmarking universal single-copy orthologue
LBA: Long-branch attraction
NGS: next-generation sequencing
LCWGS: low-coverage whole-genome sequencing
AHE: anchored hybrid enrichment
UCE: ultraconserved element enrichment
aTRAM: automated target restricted assembly method
aUBF: Average ultrafast bootstraps
GHOST: General Heterogeneous evolution On a Single Topology
PMSF: posterior mean site frequency
MSC: multi-species coalescent
gCF: gene concordance factor
sCF: site concordance factor
ML: maximum-likelihood
BI: Bayesian inference
SH-aLRT: Shimodaira-Hasegawa-like approximate likelihood ratio test
UFBS: ultrafast bootstraps
ILS: incomplete lineage sorting
EBP: Earth BioGenome Project
HGT: horizontal gene transfer
NCBI: National center for biotechnology information
BOLD: Barcode of life data system
COI: cytochrome c oxidase subunit I
LPP: local posterior probabilities
AU: approximately the unbiased
WKH: weighted Kishino-Hasegawa
WSH: weighted Shimodaira-Hasegawa
SRH: stationary, reversible and homogeneous
PISs: parsimony informative sites.

## Supplementary Information

**Additional file 1: Table S1.** Collecting information of specimens used for phylogenomic analysis in the present study. **Table S2.** The statistics for genomes of specimens from NCBI used for phylogenomic analysis in the present study. **Table S3.** The statistics for transcriptome assembly of specimens from NCBI used for phylogenomic analysis in the present study. **Table S4.** The statistics for transcriptome or genome assembly of specimens from NCBI not used for phylogenomic analysis in the present study. **Table S5**. Summary of branch support across ML and ASTRAL analyses. **Table S6.** The summarize of tree topologies consistency supported by different models and datasets in the present study. **Table S7.** Average gCF and sCF at the different classification level shown the discordance among gene trees and species trees in different datasets.

**Additional file 2: Figure S1.** Tree inferred from a maximum likelihood analysis of the BUSCO30 dataset based on partition model in IQtree. Branch support is shown on branches with the SH-like approximate likelihood ratio test value as first number and the ultrafast bootstrap value as the second number (SH-aLRT/UFB). **Figure S2.** Tree inferred from a maximum likelihood analysis of the BUSCO30 dataset based on GHOST model in IQtree. Branch support is shown on branches with the SH-like approximate likelihood ratio test value as first number and the ultrafast bootstrap value as the second number (SH-aLRT/UFB). **Figure S3**. Tree and concordance factors inferred from a maximum likelihood analysis of the BUSCO30 dataset based on PMSF model in IQtree. Branch support is shown on branches with the SH-like approximate likelihood ratio test value, the ultrafast bootstrap value, gene concordant (gCF) and site concordant (sCF) factors as the first, second, thrid and fourth numbers, respectively (SH-aLRT/UFB/gCF/sCF). **Figure S4**. Tree inferred from a maximum likelihood analysis of the BUSCO60 dataset based on partition model in IQtree. Branch support is shown on branches with the SH-like approximate likelihood ratio test value as first number and the ultrafast bootstrap value as the second number (SH-aLRT/UFB). **Figure S5.** Tree inferred from a maximum likelihood analysis of the BUSCO60 dataset based on GHOST model in IQtree. Branch support is shown on branches with the SH-like approximate likelihood ratio test value as first number and the ultrafast bootstrap value as the second number (SH-aLRT/UFB). **Figure S6**. Tree and concordance factors inferred from a maximum likelihood analysis of the BUSCO60 dataset based on PMSF model in IQtree. Branch support is shown on branches with the SH-like approximate likelihood ratio test value, the ultrafast bootstrap value, gene concordant (gCF) and site concordant (sCF) factors as the first, second, thrid and fourth numbers, respectively (SH-aLRT/UFB/gCF/sCF). **Figure S7.** Tree inferred from a maximum likelihood analysis of the BUSCO70 dataset based on partition model in IQtree. Branch support is shown on branches with the SH-like approximate likelihood ratio test value as first number and the ultrafast bootstrap value as the second number (SH-aLRT/UFB). **Figure S8.** Tree inferred from a maximum likelihood analysis of the BUSCO70 dataset based on GHOST model in IQtree. Branch support is shown on branches with the SH-like approximate likelihood ratio test value as first number and the ultrafast bootstrap value as the second number (SH-aLRT/UFB). **Figure S9**. Tree and concordance factors inferred from a maximum likelihood analysis of the BUSCO70 dataset based on PMSF model in IQtree. Branch support is shown on branches with the SH-like approximate likelihood ratio test value, the ultrafast bootstrap value, gene concordant (gCF) and site concordant (sCF) factors as the first, second, thrid and fourth numbers, respectively (SH-aLRT/UFB/gCF/sCF). **Figure S10**. Tree and concordance factors inferred from a maximum likelihood analysis of the BUSCO80 dataset based on partition model in IQtree. Branch support is shown on branches with the SH-like approximate likelihood ratio test value, the ultrafast bootstrap value, gene concordant (gCF) and site concordant (sCF) factors as the first, second, thrid and fourth numbers, respectively (SH-aLRT/UFB/gCF/sCF). **Figure S11**. Tree inferred from a maximum likelihood analysis of the BUSCO80 dataset based on GHOST model in IQtree. Branch support is shown on branches with the SH-like approximate likelihood ratio test value as first number and the ultrafast bootstrap value as the second number (SH-aLRT/UFB). **Figure S12**. Tree inferred from a maximum likelihood analysis of the BUSCO80 dataset based on PMSF model in IQtree. Branch support is shown on branches with the SH-like approximate likelihood ratio test value as first number and the ultrafast bootstrap value as the second number (SH-aLRT/UFB). **Figure S13**. Tree inferred from a maximum likelihood analysis of the lep_busco dataset based on partition model in IQtree. Branch support is shown on branches with the SH-like approximate likelihood ratio test value as first number and the ultrafast bootstrap value as the second number (SH-aLRT/UFB). **Figure S14.** Tree inferred from a maximum likelihood analysis of the lep_busco dataset based on GHOST model in IQtree. Branch support is shown on branches with the SH-like approximate likelihood ratio test value as first number and the ultrafast bootstrap value as the second number (SH-aLRT/UFB). **Figure S15.** Tree and concordance factors inferred from a maximum likelihood analysis of the lep_busco dataset based on PMSF model in IQtree. Branch support is shown on branches with the SH-like approximate likelihood ratio test value, the ultrafast bootstrap value, gene concordant (gCF) and site concordant (sCF) factors as the first, second, thrid and fourth numbers, respectively (SH-aLRT/UFB/gCF/sCF). **Figure S16**. Tree inferred from a maximum likelihood analysis of the ditrysia_busco dataset based on partition model in IQtree. Branch support is shown on branches with the SH-like approximate likelihood ratio test value as first number and the ultrafast bootstrap value as the second number (SH-aLRT/UFB). **Figure S17**. Tree inferred from a maximum likelihood analysis of the ditrysia_busco dataset based on GHOST model in IQtree. Branch support is shown on branches with the SH-like approximate likelihood ratio test value as first number and the ultrafast bootstrap value as the second number (SH-aLRT/UFB). **Figure S18**. Tree and concordance factors inferred from a maximum likelihood analysis of the ditrysia_busco dataset based on PMSF model in IQtree. Branch support is shown on branches with the SH-like approximate likelihood ratio test value, the ultrafast bootstrap value, gene concordant (gCF) and site concordant (sCF) factors as the first, second, thrid and fourth numbers, respectively (SH-aLRT/UFB/gCF/sCF). **Figure S19**. Tree inferred from a maximum likelihood analysis of the nonditrysia_busco dataset based on partition model in IQtree. Branch support is shown on branches with the SH-like approximate likelihood ratio test value as first number and the ultrafast bootstrap value as the second number (SH-aLRT/UFB). **Figure S20**. Tree inferred from a maximum likelihood analysis of the nonditrysia_busco dataset based on GHOST model in IQtree. Branch support is shown on branches with the SH-like approximate likelihood ratio test value as first number and the ultrafast bootstrap value as the second number (SH-aLRT/UFB). **Figure S21**. Tree inferred from a maximum likelihood analysis of the nonditrysia_busco dataset based on PMSF model in IQtree. Branch support is shown on branches with the SH-like approximate likelihood ratio test value as first number and the ultrafast bootstrap value as the second number (SH-aLRT/UFB). **Figure S22.** Tree inferred from a maximum likelihood analysis of the BUSCO30_test dataset based on PMSF model in IQtree. Branch support is shown on branches with the SH-like approximate likelihood ratio test value as first number and the ultrafast bootstrap value as the second number (SH-aLRT/UFB). **Figure S23**. Tree inferred from a Bayesian inference of the BUSCO30 dataset based on CAT+GTR model in PhyloBayes. Branch support is shown on branches with Bayesian posterior probability (PP) values. Two separate chains were run for 3,160-3,443 generations, then using a burn-in of 2600, and sub-sampling every 10 trees, the bpcomp program output the final consensus tree with maxdiff = 0.857143 observed across all bipartitions, but the effective size was not exactly over 300. **Figure S24**. Tree inferred from a Bayesian inference of the lep_busco dataset based on CAT+GTR model in PhyloBayes. Branch support is shown on branches with Bayesian posterior probability (PP) values. Two separate chains were run for 1,723-1,725 generations, then using a burn-in of 1000, and sub-sampling every 10 trees, the bpcomp program output the final consensus tree with maxdiff = 0.0972222 observed across all bipartitions, but the effective size was not exactly over 300. **Figure S25**. ASTRAL analysis with concordance factors of the BUSCO30 dataset. Numbers on branches represent local posterior probabilities as calculated in ASTRAL, gene concordant (gCF) and site concordant (sCF) factors calculated in IQtree. **Figure S26**. ASTRAL analysis with concordance factors of the BUSCO60 dataset. Numbers on branches represent local posterior probabilities as calculated in ASTRAL, gene concordant (gCF) and site concordant (sCF) factors calculated in IQtree. **Figure S27**. ASTRAL analysis with concordance factors of the BUSCO70 dataset. Numbers on branches represent local posterior probabilities as calculated in ASTRAL, gene concordant (gCF) and site concordant (sCF) factors calculated in IQtree. **Figure S28**. ASTRAL analysis with concordance factors of the BUSCO80 dataset. Numbers on branches represent local posterior probabilities as calculated in ASTRAL, gene concordant (gCF) and site concordant (sCF) factors calculated in IQtree. **Figure S29.** ASTRAL analysis with concordance factors of the lep_busco dataset. Numbers on branches represent local posterior probabilities as calculated in ASTRAL, gene concordant (gCF) and site concordant (sCF) factors calculated in IQtree. **Figure S30**. ASTRAL analysis with concordance factors of the ditrysia_busco dataset. Numbers on branches represent local posterior probabilities as calculated in ASTRAL, gene concordant (gCF) and site concordant (sCF) factors calculated in IQtree. **Figure S31**. Summary of tree and concordance factors inferred from a maximum likelihood analysis of the BUSCO30, BUSCO60, BUSCO70 and BUSCO80 dataset based on PMSF model in IQtree, respectively. Branch support is shown on branches with the SH-like approximate likelihood ratio test value, the ultrafast bootstrap value, gene concordant (gCF) and site concordant (sCF) factors as the first, second, thrid and fourth numbers, respectively (SH-aLRT/UFB/gCF/sCF). Confused relationship among superfamilies was underlined by gray. Superfamilies marked by red are the common taxa with unstable position of ditrysia, and that marked by blue are dynamic extracting taxa among these four datasets. **Figure S32**. The links of gene concordant (gCF) and site concordant (sCF) factors across all nodes of the ML and ASTRAL trees. (A) Concatenated ML tree under the PMSF model of BUSCO60 dataset. (B) ASTRAL tree of BUSCO60. (C) Concatenated ML tree under the PMSF model of BUSCO70 dataset. (D) ASTRAL tree of BUSCO70. (E) Concatenated ML tree under the partition scheme of BUSCO80 dataset. (F) ASTRAL tree of BUSCO80. (G) Concatenated ML tree under the PMSF model of ditrysia_busco dataset. (H) ASTRAL tree of ditrysia_busco. The colors in the figure represent three different classification levels (superfamily, family and genus), respectively.

## Acknowledgements

The authors thank Dr. Hou-Shuai Wang (South China Agricultural University, China), Mr. Bin Chen (Hunan Agricultural University, China) and Mr. Chengqing Liao (Hunan Agricultural University, China) for their kind help in collecting the samples. This study was supported by the National Natural Science Foundation of China (31970450, 41661011, 32111540167) and China Agriculture Research System (CARS-23-C08).

## Authors’ contributions

GH, XW, LC: data collection, conceived and designed the study, methodology, supervision, funding acquisition, review and editing. QC: data collection, methodology, software, data curation, visualization, writing original draft. MD: methodology, data analysis, data curation. WW: data collection, supervision, investigation. All authors have read and agreed to the published version of the manuscript.

## Availability of data and materials

Illumina sequence reads generated in this study have been deposited at NCBI’ s short sequence read archive (SRA) under accession number XXX and genome assemblies have been deposited in GenBank (BioProject accession XXX). The samples and the voucher specimens used in this study are deposited at the Insect Museum of Hunan Agricultural University. Information on the samples can be found in Additional file 1: Table S1. All of the alignments analysed in this study can be accessed from Zenodo, DOI: XXX.

## Declarations

### Ethics approval and consent to participate

Not applicable.

### Consent for publication

Not applicable.

### Author details

^1^ College of Plant Protection, Hunan Provincial Key Laboratory for Biology and Control of Plant Diseases and Insect Pests, Hunan Agricultural University, Changsha, Hunan 410128, China. ^2^ College of Science, Qiongtai Normal University, Haikou, Hainan 571100, China. ^3^ Qiannan Polytechnic for Nationality, Duyun, Guizhou 558022, China. ^4^ Research Center for Wild Animal and Plant Resource Protection and Utilization, Qiongtai Normal University, Haikou, Hainan, 571127, China. ^5^ Guangdong Provincial Key Laboratory of Silviculture, Protection and Utilization, Guangdong Academy of Forestry, Guangzhou, Guangdong 510520, China.

